# Organization of thalamocortical structural covariance and a corresponding 3D atlas of the mouse thalamus

**DOI:** 10.1101/2022.03.10.483857

**Authors:** Yohan Yee, Jacob Ellegood, Leon French, Jason P. Lerch

## Abstract

For information from sensory organs to be processed by the brain, it is usually passed to appropriate areas of the cerebral cortex. Almost all of this information passes through the thalamus, a relay structure that reciprocally connects to the vast majority of the cortex. The thalamus facilitates this information transfer through a set of thalamocortical connections that vary in cellular structure, molecular profiles, innervation patterns, and firing rates. Additionally, corticothalamic connections allow for intracortical information transfer through the thalamus. These efferent and afferent connections between the thalamus and cortex have been the focus of many studies, and the importance of cortical connectivity in defining thalamus anatomy is demonstrated by multiple studies that parcellate the thalamus based on cortical connectivity profiles.

Here, we examine correlated morphological variation between the thalamus and cortex, or thalamocortical structural covariance. For each voxel in the thalamus as a seed, we construct a cortical structural covariance map that represents correlated cortical volume variation, and examine whether high structural covariance is observed in cortical areas that are functionally relevant to the seed. Then, using these cortical structural covariance maps as features, we subdivide the thalamus into six non-overlapping regions (clusters of voxels), and assess whether cortical structural covariance is associated with cortical connectivity that specifically originates from these regions.

We show that cortical structural covariance is high in areas of the cortex that are functionally related to the seed voxel, cortical structural covariance varies along cortical depth, and sharp transitions in cortical structural covariance profiles are observed when varying seed locations in the thalamus. Subdividing the thalamus based on structural covariance, we additionally demonstrate that the six thalamic clusters of voxels stratify cortical structural covariance along the dorsal-ventral, medial-lateral, and anterior-posterior axes. These cluster-associated structural covariance patterns are prominently detected in cortical regions innervated by fibers projecting out of their related thalamic subdivisions.

Together, these results advance our understanding of how the thalamus and the cortex couple in their volumes. Our results indicate that these volume correlations reflect functional organization and structural connectivity, and further provides a novel segmentation of the mouse thalamus that can be used to examine thalamic structural variation and thalamocortical structural covariation in disease models.

## **1.** Introduction

### 1.1. The thalamus and its connections

Situated below the neocortex, the thalamus is a brain structure composed of a heterogeneous group of nuclei that, when viewed together, sends projections to all parts of the cortex (Guillery & Sherman, 2002). These nuclei reciprocally connect to the cortex (Harris et al., 2019); they relay signals from sensory organs and further modulate cortical activity through the action of intrathalamic GABAergic interneurons as part of feedback loops (Guillery & Sherman, 2002; Sherman & Guillery, 2002; Sherman, 2016). Thalamic organization may be conceptualized along various ontological axes and categories. Historic examinations of the thalamus have shown the presence of cytoarchitectonically different nuclei (Jones, 1985; Guillery & Sherman, 2002), which may be further understood through their functional roles that are derived from the cortical areas to which they primarily project (Passingham et al., 2002). These thalamic nuclei may be classified as first-order or higher-order based on the sources of information that are relayed to the cortex (Guillery & Sherman, 2002; Sherman, 2007), as part of a hierarchy of structures segmented based on multimodal imaging data that includes viral tracing (Wang et al., 2020), or through a core-matrix framework that divides thalamocortical projection neurons that emanate from these nuclei into two classes (Jones, 1998). In this latter framework, thalamocortical projections are thought to be part of either a matrix of Calbindin expressing neurons present throughout the thalamus that diffusely project to supragranular layers of many areas of the cortex, or a core set of Parvalbumin expressing neurons that primarily emerge from specific nuclei and project to the granular layers of specific areas (Jones, 1998). In all of these cases, cortical relationships are prominent in defining thalamic axes and subdivisions.

Traditional delineations of thalamic nuclei have relied on observations or histology of dissected brains, with the earliest known clear delineations attributed to Burdach (Jones, 1985). The acceptance of the role of the thalamus as an information relay to the cortex, along with the ability to infer connectivity (both structural and functional) using MRI, has yielded further insights into the connectivity-related organization of the thalamus. Using diffusion tensor imaging (DTI) and probabilistic tractography, Behrens et al. (2003) demonstrated that traditional parcellations of the thalamus can be determined from the connectivity patterns of thalamic voxels to the cortex. Subsequent MRI studies on thalamocortical connectivity have examined the relationship of these segmentations with function and cytoarchitecture (Johansen-Berg et al., 2005), assessed segmentation reproducibility (Traynor et al., 2010), examined segmentations constructed using multivariate approaches (O’Muircheartaigh et al., 2011) and statistical feature maps (Lambert et al., 2017), and applied similar methods to diffusion data to accurately target thalamic nuclei in deep brain stimulation (Middlebrooks et al., 2018). Beyond structural connectivity, other common approaches to segmenting the thalamus involve using functional relationships between the thalamus and the cortex (Eickhoff et al., 2018). Similar to DTI, resting-state functional MRI (rs-fMRI) has been used to parcellate the thalamus in young adults (Zhang et al., 2008; Yuan et al., 2016) and infants (Toulmin et al., 2015), and segmentation algorithms have been modified to capture corresponding cortical parcellations Long et al. (2021) and dynamical states (Ji et al., 2016). Recently, Müller et al. (2020) examined the core-matrix framework by dividing the thalamus based on the relative expression of Parvalbumin and Calbindin. In doing so, they showed that thalamocortical functional con-nectivity (associated with these gradients of thalamic gene expression) reflect cortical functional gradients (Müller et al., 2020).

### 1.2. Thalamocortical structural covariance

Like structural and functional connectivity, another approach to describing interregional relationships between a pair of brain regions using MRI is *structural covariance*. Structural covariance can be computed by correlating the structural properties (such as volumes) of pairs of brain regions across a group of individuals (Lerch et al., 2006; Alexander-Bloch et al., 2013a; Evans, 2013). Previous works have shown that structural covariance is associated with both structural (Lerch et al., 2006; Gong et al., 2012; Yee et al., 2018) and functional (Segall et al., 2012) connectivity, and also captures modes of coordinated growth (Alexander-Bloch et al., 2013b), potentially due to the spatially-coordinated expression of genes (Romero-Garcia et al., 2018; Yee et al., 2018). Similar to structural and functional connectivity, seed-based structural covariance maps have been used to parcellate structures including the striatum (Cohen et al., 2008), insula Kelly et al. (2012), and hippocampus Plachti et al. (2019).

At present, however, a comprehensive description of thalamocortical structural covariance is missing, and despite the thalamus being the subject of many connectivity-based parcellations, it has—to the best of our knowledge—not been segmented using structural covariance. A description of thalamocortical structural covariance would provide a deeper understanding of how the substructures within the thalamus and cortex grow together due to putative mechanisms that include coordinated neurodevelopment and plasticity. Consequently, disruptions in processes that affect cortical structure (for example, in neurodevelopmental disorders) may also result in correlated aberrant thalamic structural changes that structural covariance-based parcellations would be well suited to capture.

In this study, we use high-resolution mouse MRI to image thalamocortical structural covariance in the mouse brain in voxel-level detail. We leverage the wealth of openly available datasets on mouse brain connectivity (Oh et al., 2014) to understand the biological correlates of thalamocortical structural covariance. We sought to 1) examine the organization of thalamocortical structural covariance patterns in the mouse brain, 2) use these patterns to subdivide the mouse thalamus, and 3) understand the role of connectivity in relation to these subdivisions.

We achieve this by constructing voxelwise structural covariance maps in the cerebral cortex for each voxel in the thalamus as a seed. In conjunction with external tract-tracing data, we further describe the association between thalamocortical structural covariance and structural connectivity. Our results provide both an insight into the organizational principles of thalamocortical structural covariance, and also a data-driven atlas of the thalamus that would provide a matched filter to optimize detection of thalamocortical structural covariance in future mouse MRI studies.

## 2. Methods

### 2.1. Overview

In this study, we use a population of 154 healthy control mice to compute thalamocortical structural covariance^1^. We use the Fisher-transformed Pearson correlation coefficient, computed between the local volumes of seed voxels in the thalamus and target voxels in the cortex, to measure structural covariance. Restricting our analysis to a single hemisphere for computational reasons, we compute ipsilateral structural covariance in the cortex (7643 voxels at 200*µ*m isotropic resolution) using every voxel in the left thalamus as a seed (80456 voxels at 50 *µ*m isotropic resolution).

The aforementioned set of structural covariance maps (a 80456 × 7643 matrix) were examined in relation to thalamocortical structural connectivity and clustered to identify clusters of thalamic voxels that were similar in their cortical structural covariance patterns. **Figure 1** provides a graphical overview of the procedure used to compute thalamocortical structural covariance.

**Figure 1:**
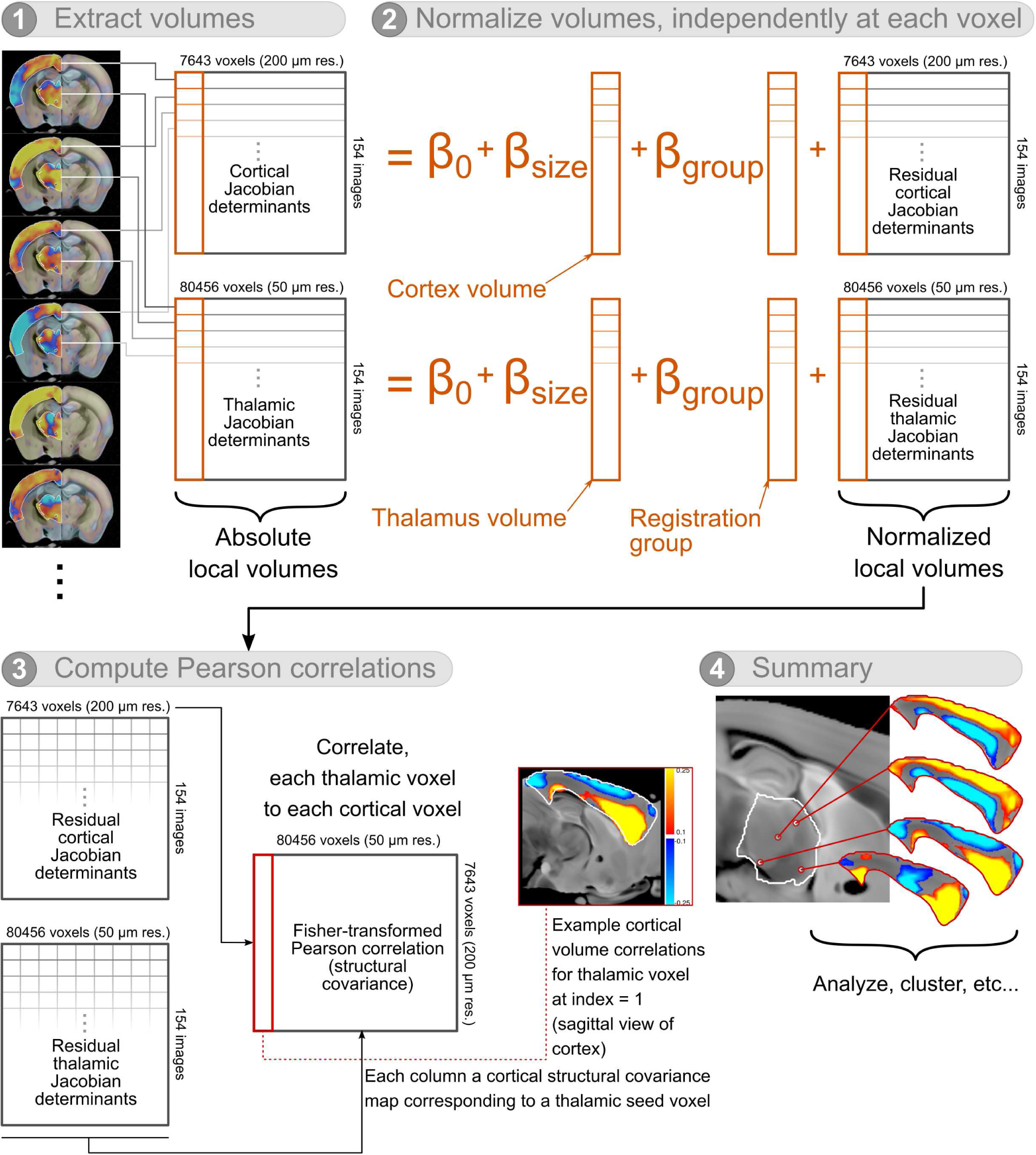
A graphical overview of the procedure used in computing thalamocortical structural covariance. For visualization purposes, cortical maps of structural covariance are upsampled to 50 *µ*m. Also for ease of visualization, the structural covariance matrix is shown transposed; in the analysis, it is treated as a 80456 × 7643 matrix.

### 2.2. Mouse population and imaging

All structural covariance maps are computed over a population of 154 wildtype mice that are controls from a larger dataset, a smaller part of which is described by Ellegood et al. (2015). Images from this dataset were selected such that mice are as identical as possible, so that as little variation and subsequent covariation can be attributed to variation in age, strain, environmental conditions, or scan sequence differences. The obtained sample size is well above the sixty mice Pagani et al. (2016) recommended for computing structural covariance.

In this case, we pooled together control mouse brain images from eight different datasets; all images were of mice that were all young adults (P56-84), of both sexes, and of either the C57Bl/6J (“B6J”) or C57Bl/6N (“B6N”) substrains that make up the widely used C57Bl/6 inbred strain. Images were acquired *ex vivo* (with brains left within the skull) in parallel, sixteen at a time, as described by previous studies (Dazai et al., 2011; Lerch et al., 2011; Nieman et al., 2018). All images were acquired on the same 7T Varian MR scanner using the same *T*_2_-weighted 3D fast spin echo sequence that resulted in whole brain images at an isotropic resolution of 40 *µ*m after 14 hours of scanning (Spencer Noakes et al., 2017). **Table 1** describes the population of mice.

**Table 1:** A description of the imaging datasets used to construct thalamocortical structural covariance maps, including mouse numbers, strain, sex, age.

| Dataset | Sample size | Sex | Strain | Age | Scanner | Scan sequence | Scan resolution |
| --- | --- | --- | --- | --- | --- | --- | --- |
| 1 | 35 | Both (13F) | C57Bl/6N | P58-64 | 7T Varian | $T_2w$ FSE 3D | 40 $\mu m$ |
| 2 | 12 | M | C57Bl/6J | P56 | 7T Varian | $T_2w$ FSE 3D | 40 $\mu m$ |
| 3 | 10 | M | C57Bl/6J | P56 | 7T Varian | $T_2w$ FSE 3D | 40 $\mu m$ |
| 4 | 11 | Both (7F) | C57Bl/6N | P77 | 7T Varian | $T_2w$ FSE 3D | 40 $\mu m$ |
| 5 | 23 | Both (9F) | C57Bl/6N | P84 | 7T Varian | $T_2w$ FSE 3D | 40 $\mu m$ |
| 6 | 16 | Both (10F) | C57Bl/6J | P56-70 | 7T Varian | $T_2w$ FSE 3D | 40 $\mu m$ |
| 7 | 40 | Both (20F) | C57Bl/6J | P63-70 | 7T Varian | $T_2w$ FSE 3D | 40 $\mu m$ |
| 8 | 7 | M | C57Bl/6J | P70-84 | 7T Varian | $T_2w$ FSE 3D | 40 $\mu m$ |
| Summary | 154 | Both | C57Bl/6 | P56-84 | 7T Varian | $T_2w$ FSE 3D | 40 $\mu m$ |

### 2.3. Registration and volumes

Whole brain MR images, corrected for *B*_0_ distortions (Nieman et al., 2018) and intensity non-uniformities (Sled et al., 1998), were registered together as part of a larger study (in preparation) using the Pydpiper registration framework (Friedel et al., 2014). No mice were excluded since the scans were acquired and quality-controlled as part of other experiments.

Briefly, this registration is carried out in two main levels. In the first level of registration, images within each individual dataset (eight datasets in our case) are registered together to form an unbiased study-specific average brain. This registration involves an iterative series of six-parameter linear (rotations and translations), twelve-parameter affine (rotations, translations, scales, and shears), and nonlinear alignments, where in each step individual images are aligned to the previous step’s average. Linear alignments are computed using minctracc (Collins et al., 1994) and nonlinear alignment of images are performed using ANTs (Avants et al., 2008, 2011). Outputs of this first level of registrations include Jacobian determinant images—associated with each input image and defined over the study average—that encode local volume information. In the second level of the pipeline, the first-level dataset averages are registered together in the same process described above. Jacobian determinants associated with each image, mapped onto the global second-level average constructed from all studies, are obtained by concatenating first-level and second-level transformations together.

Finally, to allow for comparisons with the Allen Institute’s resources that are defined within their Common Coordinate Framework (version 3; CCFv3) (Wang et al., 2020), we aligned our global template to their CCFv3 template (an average of *>*1600 autofluorescence images obtained via serial two-photon microscopy). Registration was performed using ANTs and each Jacobian determinant image was further resampled to CCFv3 space; thus, for each mouse, there exists a Jacobian determinant map defined over the CCFv3 template that describes local volumes at each voxel. This MRI to CCFv3 registration was visually inspected and confirmed to be of high quality; this can be verified in various subsequent figures (e.g. **Figure S1a**) that display the co-registered data across hemispheres. All analyses were carried out in CCFv3 space.

### 2.4. Cortex and thalamus definitions

In this study, voxels within the thalamus were considered as seeds; for each seed voxel, structural covariance (described below) was computed to every voxel in the cortex. We used the Allen Institute’s Mouse Brain Atlas (AMBA) (Wang et al., 2020) definitions of the thalamus and cortex as regions in which our analyses are restricted. Note: in these definitions, the subthalamic nucleus, zona incerta, and fields of Forel (which some define as the “ventral thalamus”) are part of the hypothalamus and not considered here; the corresponding “dorsal thalamus”—the part which contains cortical efferent and afferent fibers and includes the thalamic reticular nucleus—are simply referred to here as the thalamus (in line with the AMBA definitions) and form the basis of this study. We obtained structure masks from their download server at 50 *µ*m resolution (structure ID = 315, corresponding to the Isocortex, and structure ID = 549, corresponding to the thalamus). These structures are defined over the Allen Institute’s CCFv3 template, to which our MR data were aligned.

For computational reasons, we computed cortical structural covariance at a 200 *µ*m resolution. To be clear, this down-sampling was carried out only for cortical data; the sizes of seed voxels within the thalamus were kept at 50 *µ*m along each axis, and subsequent analyses and segmentations within the thalamus were carried out at this higher resolution. Furthermore, also for computational efficiency, thalamic and cortical structure masks were further intersected with a left hemisphere mask, and the resulting structural covariance computations and analyses were carried out in this single hemisphere. We note that a consistent finding across multiple studies is that seed-based structural covariance maps are generally bilateral (i.e. homotopic regions tend to have similar structural covariance to seed (Mechelli et al., 2005; Pagani et al., 2016; Yee et al., 2018), reflecting the bilateral patterns of transcriptomic similarity, a major correlate of structural covariance (Yee et al., 2018)). Thus, including both hemispheres would be redundant and would likely give similar results to our findings below.

**Figure S1a** shows the 50 *µ*m thalamic mask under which each voxel was used as a seed, and the 200 *µ*m cortical mask within which cortical structural covariance was computed. At these resolutions, there were 80456 voxels within the thalamus (left hemisphere) mask and 7643 voxels within the cortex (left hemisphere). Distributions of cortical, thalamic, and whole brain volumes (**Figure S1b**) were further examined as a measure of registration and volume computation quality, and were found to be within expected ranges of volumes.

### 2.5. Structural covariance computation

In this study, we use volume as the structural property from which structural covariance is computed. As measures of local volume, we use the log-transformed Jacobian determinant, and for each voxel, further regress out 1) the effect of dataset (the first column in **Table 1**, a categorical variable), and 2) the absolute volume of the left hemisphere isocortex or left hemisphere thalamus, depending on the location of the voxel.

The remaining residuals describe volume deviations relative to the size of its parent structure (isocortex or thalamus) for each subject, and from these data, we compute the Pearson correlation coefficient (i.e. normalized covariance) between values at the seed and target voxels. Correlation coefficients are further variance-stabilized via the Fisher transformation. Guided by the results of Yee et al. (2018) that suggests that transcriptomic similarity is an important contributor to the structural covariance signal, the subsequent analyses described are performed using correlation data in which transcriptomic similarity is not regressed out. We did however repeat all analyses after regressing out distance and transcriptomic similarity and found a negligible difference in the overall results; for brevity, these are not presented here.

The resulting set of 80456 7643-element vectors of Fisher-transformed correlation coefficients—each of which describes cortical volume correlations to a given thalamic seed voxel—forms our definitions of structural covariance. Al-though the Pearson correlation is symmetric and the resulting matrix of correlations could equivalently be described as representing *thalamocortical* or *corticothalamic* structural covariance, we work with thalamic voxels as seeds and therefore refer to this structural covariance as *thalamocortical* from here on.

### 2.6. Statistical analyses

#### 2.6.1. Outline

The 80456 × 7643 structural covariance matrix formed the basis of a variety of analyses. We examined:

- whether cortical structural covariance maps (row vectors drawn from this matrix) displayed high values in regions known to be structurally and functionally connected with specific thalamic seed voxels,
- whether cortical structural covariance varied as a function of cortical depth,
- whether thalamic voxels could be clustered based on cortical structural covariance, and
- whether the cortical structural covariance components associated with each thalamic cluster could be explained by cluster-specific thalamocortical wiring.

#### 2.6.2. External datasets used

In this analysis, we use two different datasets from the Allen Institute: the Allen Mouse Brain Atlas’s (AMBA) Common Coordinate Framework (Wang et al., 2020) that defines cortical areas and nuclei within the thalamus, and the Allen Mouse Brain Connectivity Atlas (AMBCA) (Oh et al., 2014; Harris et al., 2019) that provides information on thalamocortical structural connectivity.

*Allen Mouse Brain Atlas.* The Allen Mouse Brain Atlas (AMBA) parcellates the mouse brain in a hierarchical manner. Segmentations, defined on the Common Coordinate Framework version 3 (CCFv3) template, were downloaded at 25 *µ*m isotropic resolution and resampled to 50 *µ*m resolution at which the analyses within the thalamus were done. As a node within a larger tree of structures across the entire brain, the thalamus begins at the fifth level of this tree, is further subdivided into two major subdivisions that label voxels within as either sensory-motor cortex related or polymodal association cortex related (sixth level), and contains child structures down to the ninth level.

*Allen Mouse Brain Connectivity Atlas.* The Allen Mouse Brain Connectivity Atlas (AMBCA) consists of a series of viral tracing experiments in which a recombinant adeno-associated viral (rAAV) tracer expressing a fluorescent protein was injected into various regions of the mouse brain. Injections were in both wildtype mice and Cre-lines that targeted specific cell types. These tracers travel anterograde and do not cross synapses. Images of the fluorescent tracer were obtained via serial two-photon microscopy, and further processed by the Allen Institute and aligned to the CCFv3 template.

Among these processed images are “projection density” and “injection fraction” images that were downloaded at 50 *µ*m resolution; these define the proportion of voxels that express a tracer signal at the original imaging resolution and the proportion of voxels at the original imaging resolution that are part of the manually labeled injection site.

#### 2.6.3. Cortical structural covariance and functional systems

To assess whether thalamocortical structural covariance reflected functional systems, we focused on four thalamic nuclei that have been well characterized in their functional roles and their cortical connectivity profiles:

- lateral geniculate nucleus (visual system),
- medial geniculate nucleus (auditory system),
- ventral anterior/ventral lateral nuclei (together grouped as the ventral anterior-lateral nucleus; motor system), and
- ventral posteromedial nucleus (somatosensory system).

We used the AMBA segmentations of these regions and chose the centroid voxel of these regions (structure abbreviations: LGd_co, MGv, VAL, VPM) as a seed for which cortical structural covariance maps were generated. These cortical covariance maps were visually examined in relation to 1) AMBA cortical parcellations of relevant areas (e.g. the visual cortex for the LGd_co structural covariance map), and 2) tracer projections targets for a representative neuronal tracer injection at the location of the seed.

Structural covariance maps for each of the four seeds and related cortical data were visualized on a flattened representation of the cortex (Harris et al., 2019; Wang et al., 2020).

#### 2.6.4. Cortical structural covariance and depth

To examine the dependence of cortical structural covariance on cortical depth, we first obtained a depth map of the cortex that was constructed by the Allen Institute (Wang et al., 2020) by solving Laplace’s equation within a cortical mask with appropriate boundary conditions (Jones et al., 2000; Lerch et al., 2008; Lee et al., 2011). Within this depth map, each voxel within the cortex was given a value that ranged between 0 (most superficial part of the cortex) and 1 (deepest part of the cortex). For each seed-based cortical structural covariance map, our goal was to find the depth at which highest structural covariance is observed. One option would be to directly determine the value of the depth map at the point in the cortex where highest structural covariance is observed. Rather than choosing a single point that might be a noisy estimate of true depth, we chose to average over multiple voxels with high structural covariance, and thus computed the mean cortical depth of the top 100 voxels (at 200 *µ*m resolution) displaying highest structural covariance. The choice of 100 was arbitrary. Thus, for each thalamic voxel as a seed, a depth value that characterized the location of highest positive cortical structural covariance was obtained.

To understand the major modes of cortical depths at which high structural covariance is seen, a mixture model was fitted onto the resulting distribution of depths using the mclust package (Scrucca et al., 2016) in R (R Core Team, 2020). The number of components was chosen based on the point at which the *BIC* plateaus when plotted as a function of number of components, assuming unequal variances.

#### 2.6.5. Clustering of thalamic voxels

Cortical structural covariance maps were generated for each of the 80456 voxels in the thalamus; these were clustered together to 1) identify voxels in the thalamus that share similar cortical structural covariance patterns and 2) determine what these common cortical structural covariance “components” (average cortical structural covariance for voxels within each cluster) might look like. Given the large number of observations of a high dimensional vector, we opted to use *k*-means clustering for computational efficiency. Clustering was carried out in Python using the scikit-learn’s (Pedregosa et al., 2011) implementation of the Elkan algorithm (Elkan, 2003) (sklearn.cluster.KMeans). In our purely covariance-driven approach, no explicit spatial information of seed voxel location is used for clustering. Parameters used are described in the clustering class instantiation call below:

KMeans(n_clusters=n_clusters,

n_init=10, max_iter=500,

n_jobs=5, algorithm=”elkan”, precompute_distances=False,

verbose=1)

Optimal number of clusters was determined by constructing a *scree* plot that plots the ratio of the total within-cluster sum of squares distance and between-cluster sum of squares distance as a function of number of clusters. The number of clusters was chosen as the point at which little change in this ratio is observed as the number of clusters is increased, i.e. the “elbow” in this curve.

The resulting clustering of thalamic voxels provided a set of segmentations that were compared to AMBA segmentations of the thalamus. For each pair of regions formed by pairing each structural covariance-based cluster of voxels and each AMBA segmentation of a thalamic region, we computed the binary overlap. Considering the AMBA segmentation as “ground truth”, we determined the sensitivity (proportion of AMBA structure that contains voxels within the thalamic cluster), positive predictive value (proportion of thalamic cluster that contains voxels within the AMBA structure), and Dice coefficient (the harmonic mean of the sensitivity and positive predictive value, also known as the *F*_1_ score). To compute the probability that these values could arise by chance (i.e., the *p*-values), a permutation test was used in which cluster assignments were permuted 10000 times. Subsequent *p*-values were adjusted to account for the multiple pairwise tests using the Bonferroni correction, thereby controlling for the family-wise error rate.

#### 2.6.6. Relationship to connectivity

Given the association between structural covariance and structural connectivity (Gong et al., 2012; Yee et al., 2018), we asked if the cortical structural covariance maps associated with each cluster of thalamic voxels could be explained by thalamocortical structural connectivity specific to the thalamic cluster. For each thalamic cluster, we first looked for which AMBCA experiments had injection sites that overlapped with voxels within the cluster. Viral tracing experiments were mapped to a cluster if at least half of the binarized injection fraction image (binarized at 0.9) contained voxels within that cluster. Subsequently, we merged all cluster-related tracer experiments by taking the maximum projection density at each voxel. Thus, for every cluster, we produce a single image that describes the tracts that project out of the thalamic cluster.

As a measure of association between cortical structural covariance and the merged tracer image, we compute the sensitivity of high positive and negative structural covariance (binarized above and below structural values of 0.1 and −0.1 respectively) to this merged tracer projection density (binarized at a projection density value of 0.1). To test whether cluster-specific cortical structural covariance is associated with connectivity, we use a permutation test in which we sample a random set of tracer experiments in which the injection site overlaps any thalamic cluster, keeping the number of sampled experiments equal to the number of overlapping injection sites in the original cluster. We used 5000 permutations, considered both tails of this permutation distribution in computing *p*-values, and corrected *p*-values for multiple comparisons using the Benjamini-Hochberg FDR method (Benjamini & Hochberg, 1995).

### 2.7. Data and code availability

MR imaging data will be made available through the Ontario Brain Institute’s central database, Brain-CODE. These data are part of a broader study, part of which is already available from Brain-CODE. Connectivity and reference atlas data from the Allen Institute are openly available on brain-map.org.

Open-source code is available on GitHub at: http://github.com/yohanyee/project-subdividing-the-thalamus. All packages and toolkits used are freely available, including the minc-toolkit (Vincent et al., 2016), RMINC (Lerch et al., 2017), and GNU Parallel (Tange, 2011).

## 3. Results

### 3.1. Thalamocortical structural covariance reflects sensory function

*Thalamocortical structural covariance distinguishes visual from auditory networks.* For a seed voxel within the lateral geniculate nucleus (LGd_co), a structure that relays signals to the visual cortex as part of the visual system (Hollander, 1972; Hendrickson et al., 1978), high cortical structural covariance is observed in the visual cortex (**Figure 2a**, top row, second column), consistent with size covariation reported in humans (Andrews et al., 1997). Similarly, for a seed voxel in the medial geniculate nucleus (MGv), a thalamic nucleus that relays signals to the primary auditory cortex (Hashikawa et al., 1995; Molinari et al., 1995), high structural covariance in the auditory cortex is observed (**Figure 2a**, top row, third column). Contrasting these sensory systems by taking the difference demonstrates that structural covariance maps can be used to separate functionally distinct areas of the cortex (**Figure 2a**, top row, rightmost column). Together, these results suggest that thalamocortical structural covariance reflects the functional organization of the cortex. While structural covariance might be sensitive to function, we note that it is not specific to cortical regions canonically associated with each function. High structural covariance is also present in other regions; in the case of the lateral geniculate nucleus, high structural covariance is also observed in motor and somatosensory areas, while for the medial geniculate nucleus, high structural covariance is also observed in the agranular insula and anterior cingulate areas.

**Figure 2:**
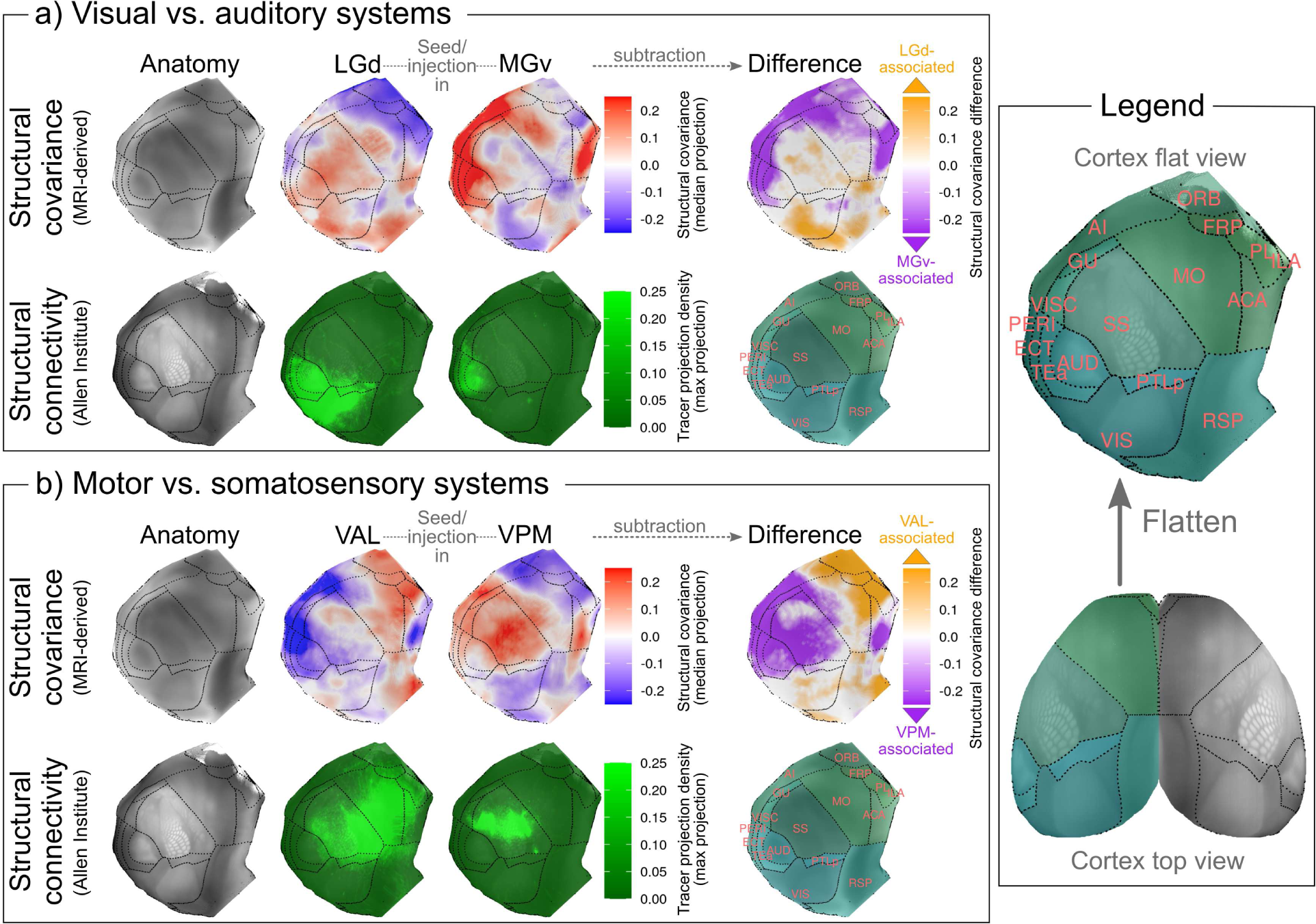
Seed-based cortical structural covariance maps, corresponding to four functionally-relevant seed voxels in: the dorsal lateral geniculate nucleus (LGd; visual function), the ventral medial geniculate nucleus (MGv; auditory function), the ventral anterior-lateral complex (VAL; motor function), and ventral posteromedial nucleus (VPM; somatosensory function). For each of the four seeds, structural covariance and connectivity data are projected onto a 2D representation of the flattened cortex (also depicted within the legend on the right, see Harris et al. (2019) and Wang et al. (2020) for more details). In **a)**, visual and auditory function-related structural covariance maps are compared and contrasted to each other; in **b)**, the same is done for motor and somatosensory function-related structural covariance. Within each subfigure, images in the top row correspond to data obtained through MR imaging and are projected on the cortical surface by taking the median value along cortical streamlines; specifically, shown from left to right are anatomy as seen through MRI, structural covariance maps for seeds in each of the two thalamic structures being compared, and their resulting difference in structural covariance. Differences in cortical structural covariance between the two functional systems highlight the spatial specificity of thalamocortical structural covariance. Note that the scale bars associated with structural covariance are clamped at 0.25 to allow for visual contrast and enable comparisons across different seed regions; there exist structural covariance values that are greater than 0.25 and less than −0.25. Specifically, for each of the four seed regions LGd, MGv, VAL, and VPM, the proportions of structural covariance that lie outside the clamped scale are 0.21%, 5.52%, 3.12%, and 0.24% respectively. In the bottom row, images correspond to reference data from the Allen Institute and are projected on the cortical surface by taking the maximum value along cortical streamlines; shown from left to right are anatomy within the CCFv3 template, viral tracer projection density for injections in each of the two thalamic structures being compared, and a surface annotated with cortical structure acronyms from the Allen Mouse Brain Atlas (AMBA). AMBA annotations are FRP: Frontal pole; MO: Somatomotor areas; SS: Somatosensory areas; GU: Gustatory areas; VISC: Visceral areas; AUD: Auditory areas; VIS: Visual areas; ACA: Anterior cingulate area; PL: Prelimbic area; ILA: Infralimbic area; ORB: Orbital area; AI: Agranular insular area; RSP: Retrosplenial area; PTLp: Posterior parietal association areas; TEa: Temporal association areas; PERI: Perirhinal area; ECT: Ectorhinal area.

To compare structural covariance to structural connectivity, we picked representative tracer experiments for an injection in the lateral (AMBCA experiment ID: 298003295; **Figure 2a**, bottom row, second column) and medial (experiment ID: 180520257; **Figure 2a**, bottom row, third column) geniculate nuclei. For the LGd as seed, we observed some overlap between cortical structural covariance and connectivity, particularly in posterior visual areas. Of note, LGd-related tracer signal and cortical structural covariance are observed in parts of the auditory area too. In the case of structural covariance to the MGv, this sensitivity to connectivity was far more convincing, with high structural covariance being observed in areas of the cortex connected to the seed. We note that in this case—as with our observations on function described above— structural covariance appears to be sensitive but not specific to structural connectivity. High structural covariance is also present in unconnected areas; while this could be due to the methodology (i.e. the tracers may not capture the true extent of thalamocortical connectivity), this is consistent with prior findings that structural covariance networks are more extensive than structural wiring (Gong et al., 2012; Yee et al., 2018).

In **Figure 2**, cortical structural covariance and connectivity are projected onto a flattened representation of the cortex so that the entire spatial extent of these signals can be seen. We observed however that cortical structural covariance also varied by depth (analyzed in detail below), and therefore provide views of the same data on slices of a brain volume (**Figure S2a**) where this can be seen.

*Thalamocortical structural covariance distinguishes motor from somatosensory networks.* The ventral nuclear group of the thalamus contains the ventral anterior nucleus (VA), the ventral lateral nucleus (VL), and the ventral posterior nucleus (VP), the latter of which consists of the ventral posteromedial nucleus (VPM) and ventral posterolateral nucleus (VPL). As relay nuclei, the VA and VL facilitate the flow of motor information to the primary and secondary motor cortex (Jones et al., 1979), while the VPM connects the thalamus to the postcentral gyrus, relaying information to the somatosensory cortex (Jones & Powell, 1970). In the AMBA ontology, the VA and VL are grouped together as the ventral anterior-lateral complex of the thalamus (VAL). Here, we examine cortical structural covariance for seeds in the VAL and VPM, thereby comparing the motor and somatosensory systems.

For structural covariance to the VAL, we observed that high structural covariance was present in parts of the motor cortex (**Figure 2b**, top row, second column). Specifically, structural covariance is observed in the secondary motor cortex and deeper layers of the posterior primary motor cortex (**Figure S2b**, left column). In contrast to the VAL, structural covariance to the VPM is seen in somatosensory areas (**Figure 2b**, top row, third column), particularly in superficial layers of the primary somatosensory cortex (**Figure S2b**, middle column). Contrasting these maps by computing the difference shows that structural covariance separates the motor cortex from somatosensory areas (**Figure 2b**, top row, rightmost column), and that this separation is better in the association (i.e. secondary) as opposed to primary areas due to structural covariance signals overlapping along the depth axis in the latter (**Figure S2b**, right column).

Cortical structural covariance to the VAL is generally observed in cortical regions that are connected to the VAL (AMBCA experiment ID: 113884251; **Figure 2b**, bottom row, second column); these appear to be medial areas of the frontal cortex. Interestingly, these projections highlight the supplementary but not primary motor cortex, consistent with structural covariance. We note that the VAL-related tracer signal also extends to parts of the somatosensory cortex where structural covariance is not observed; this is likely because part (10%) of the injection site for this tracer experiment extends into the VPM. In contrast to the VAL, neurons from the VPM project to lateral areas that are more posterior (experiment ID: 158375425; **Figure 2b**, bottom row, third column), a pattern further reflected in the cortical structural covariance maps. *Thalamocortical structural covariance shows depth specific patterns.* Beyond localization of high structural covariance to functionally-relevant areas of the cortex, cortical structural covariance was also found to vary along the depth axis. We observed a distinct pattern of thalamocortical structural covariance in which certain thalamic voxels showed high structural covariance to superficial layers (an example for one such voxel is shown in **Figure 3a**), while others showed high structural covariance to deeper layers. To analyze this relationship with depth for all thalamic seed voxels, we summarized each cortical structural covariance map by the average depth of cortical voxels that displayed the highest positive structural covariance (top 100 voxels, **Figure 3b**).

**Figure 3:**
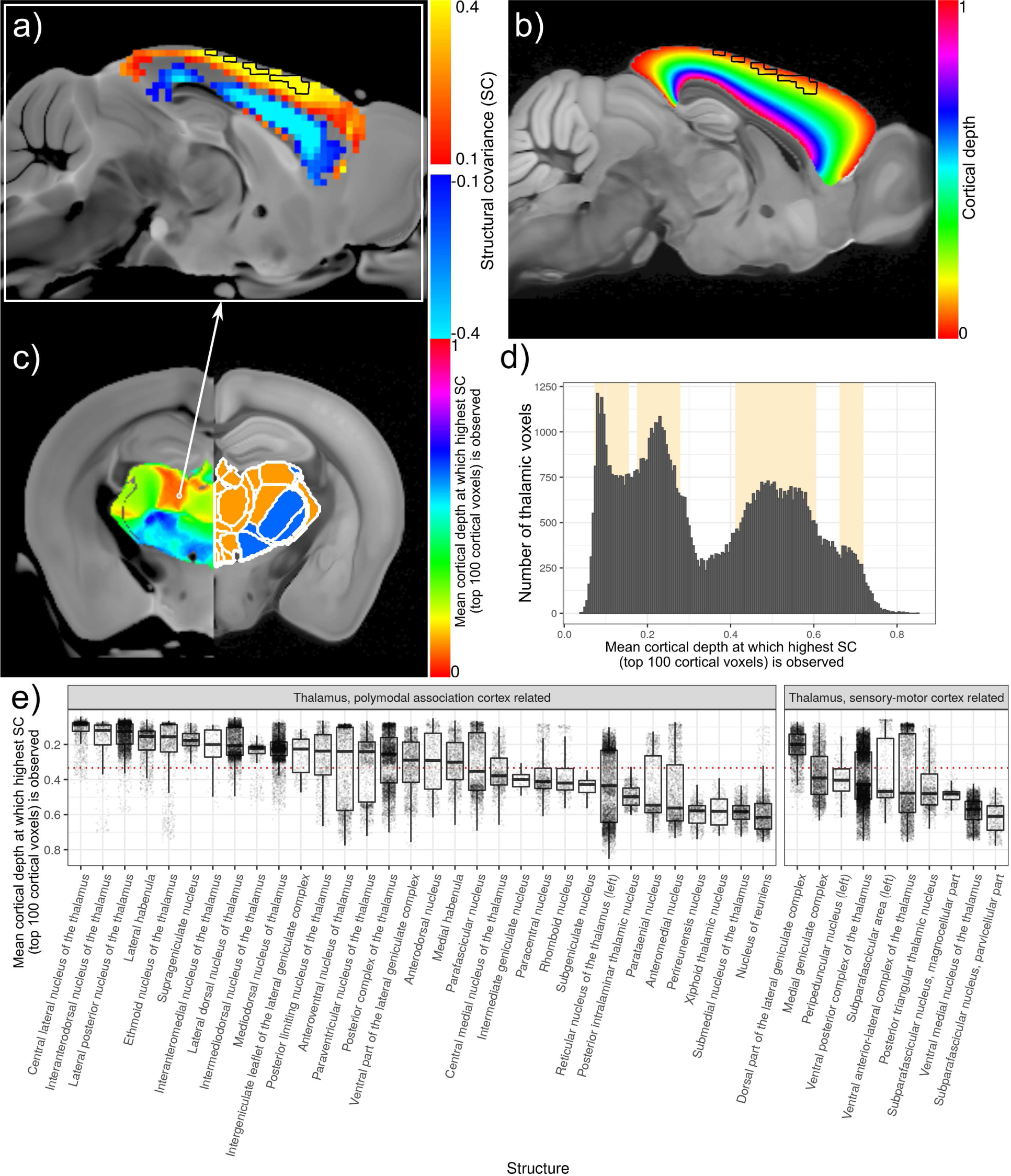
Thalamocortical structural covariance and cortical depth. In **a)**, an example of a cortical structural covariance map for a seed voxel in the central lateral nucleus of the thalamus (CL; location shown in **(c)**) is plotted on a sagittal slice with the top 100 voxels (at 200 *µ*m, ranked by structural covariance values) outlined in black. These same voxels are also highlighted in **(b)** on a map of cortical depth (scaled between 0 and 1; constructed by the Allen Institute by solving Laplace’s equation within the cortex with appropriate boundary conditions), demonstrating the quantification of the mean depth at which high positive cortical structural covariance is seen. In **c)**, this mean depth is plotted on each voxel of the thalamus based on the voxel’s cortical structural covariance map, and in **(d)**, a histogram of these values are displayed with major modes (components of a mixture model, mean ± one standard deviation) highlighted. In **e)**, this distribution is further split based on Allen Mouse Brain Atlas segmentations that are further grouped by their classification as sensory-motor cortex or polymodal association cortex-related nuclei.

We found that voxels in the dorsal thalamus tend to covary strongly with superficial voxels within the cortex, while voxels within the ventral thalamus tend to covary strongly with deeper voxels (**Figure 3c** left hemisphere); these appear to reflect the AMBA segmentations of the thalamus into sensory-motor cortex-related and polymodal association cortex-related regions (the highest subdivision of the thalamus in a hierarchy of subdivisions, **Figure 3c** right hemisphere). Across all voxels, a mixture model detected five components within the distribution of these mean depth values, two of which overlap within one standard deviation of their means (**Figure 3d**). Further examining this depth relationship by finer nuclei as defined by the AMBA, it appears that the stratification of structural covariance into distinct modes of depth is also present for seed voxels within these nuclei (**Figure 3e**). Given this variation of cortical structural covariance with cortical depth, further analyses of cortical structural covariance maps are carried out and visualized over 3D volumes, as opposed to over a 2D representation of the cortex.

We note that structural covariance is computed from volume information (Jacobian determinants) that depends on contrast obtained from *T*_2_-weighted MR images of *ex vivo* brains, and this source of contrast provides layer-related information. In the MR images that we used, layer-like structures are visible, and the five modes corresponding to the five different depths at which high structural covariance is seen (**Figure 3d**) can be mapped onto these structures (**Figure S3**). Despite the observation of five such structures that alternate in brightness, it is not completely certain whether these MR-derived structures correspond to traditional cortical layers however, since they do not always correlate with the layers observed through histology (Boretius et al., 2009), and may instead reflect myelin density (Stüber et al., 2014; Fracasso et al., 2016) or iron concentration (Fukunaga et al., 2010; Stüber et al., 2014).

### 3.2. Subdividing the thalamus

*Thalamocortical structural covariance patterns display sharp boundaries within the thalamus.* In comparing cortical structural covariance to seed voxels in the lateral geniculate and medial geniculate nuclei, we found that the cortical maps were qualitatively quite different (**Figures 2a** and **S2a**), despite the seeds being anatomically very close to each other (1.08 mm) and part of the same parent nuclei group (geniculate group). This suggests that the small nuclei within the thalamus that differ in their function may be delineated by observing transitions in their structural covariance patterns. To probe this, we swept seed voxels along various axes within the thalamus and observed sharp transitions in cortical structural covariance maps, providing additional evidence that such boundaries may be observed. Often, but not always, these boundaries roughly correspond to the divisions of thalamic nuclei. The presence of sharp boundaries suggests that thalamic voxels cluster into modules within a space that is described by distances between cortical structural covariance, and therefore a clustering approach is appropriate. We observed these boundaries for various axes such as the medial-lateral axis shown in **Figure 4**; similar results are shown in **Figure S4** for anterior-posterior, dorsal-ventral, and anteromedial-posterolateral axes, and in movies within the **Supplementary Data** for denser arrangements of seeds along these axes.

**Figure 4:**
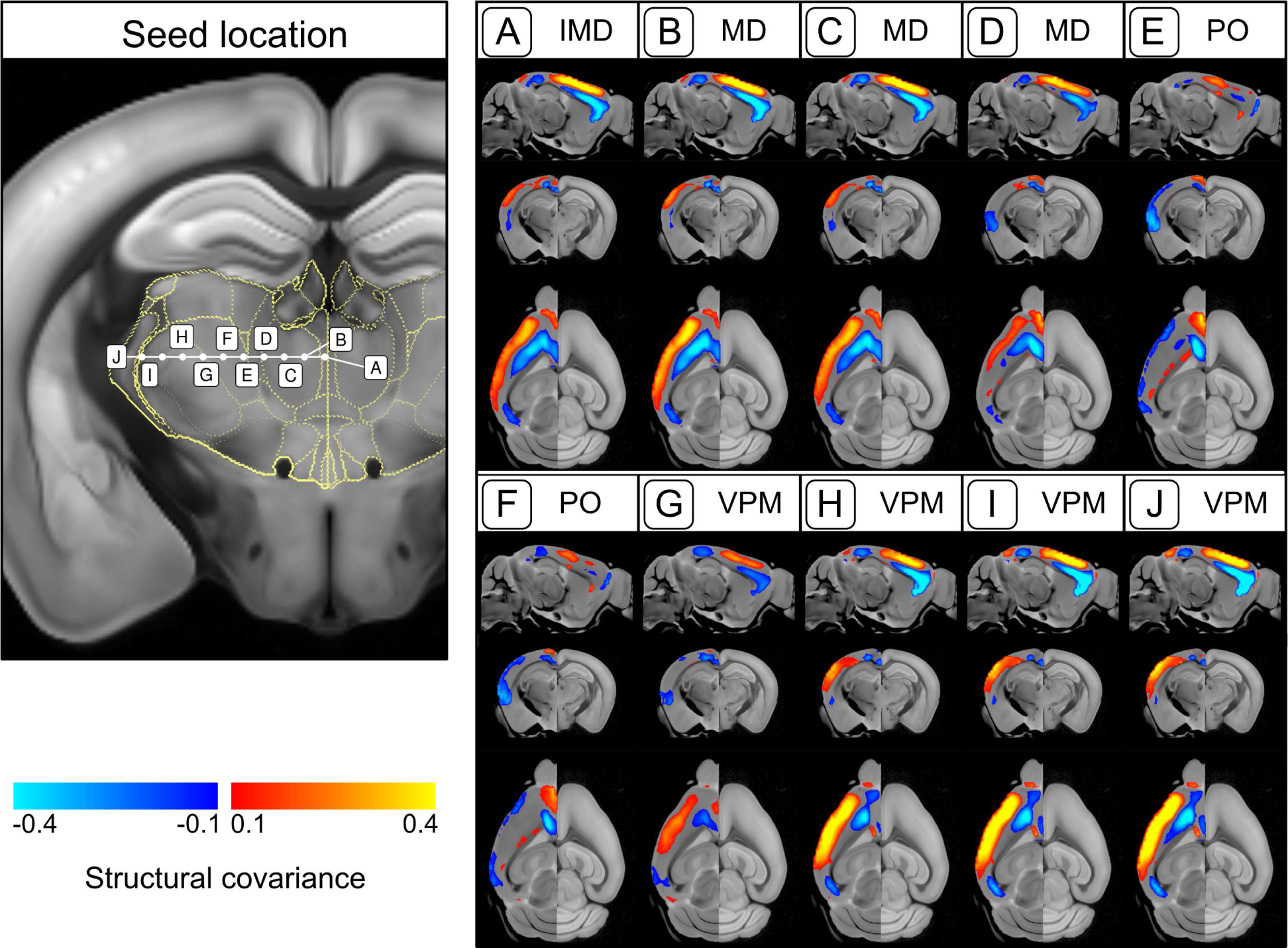
Observation of sharp transitions between cortical structural covariance of nearby seed voxels. Shown are cortical structural covariance maps for various equally-spaced seed points in the thalamus that are arranged along a medial-lateral axis (left panel). For each annotated seed point on the coronal slice on the left, a sagittal, coronal, and axial slice of the associated cortical structural covariance map (resampled to 50 *µ*m) is plotted (right panel). Labels at the top correspond to the seed point annotation, along with the thalamic nucleus (denoted by its Allen Mouse Brain Atlas acronym) that the seed lies within.

*k-means clustering of corticothalamic structural covariance.* Given the large number of thalamic voxels, we opted to use *k*-means clustering, a computationally efficient—and in this case tractable—algorithm to group thalamic voxels together based on their cortical structural covariance patterns. Subdividing the thalamus at various numbers of clusters ranging from 2 to 42, we found (based on the elbow location within a scree plot, **Figure 5a**) that six clusters described the data well.

**Figure 5:**
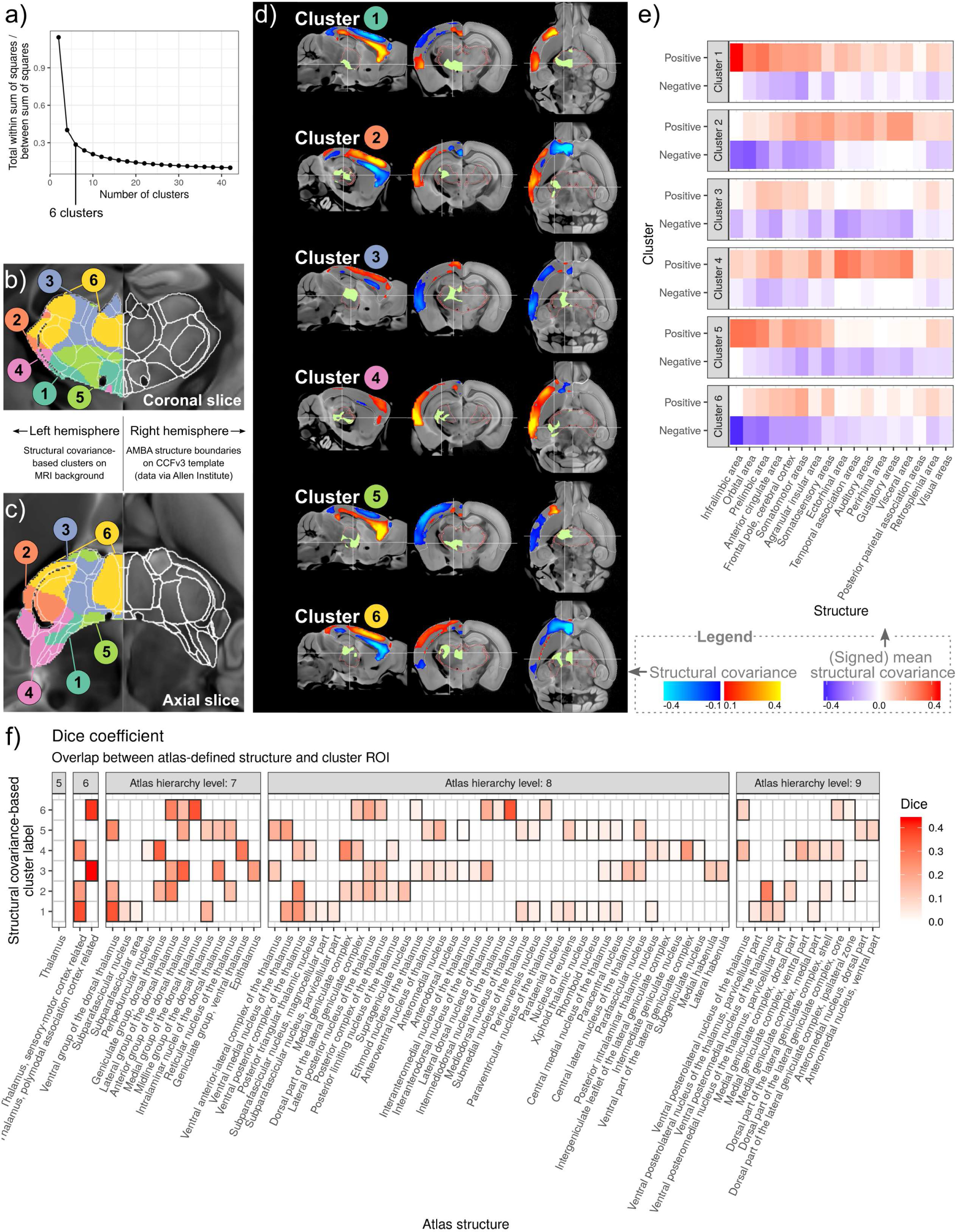
A structural covariance-based atlas of the mouse thalamus. In **a)**, a scree plot shows the ratio of the total within-cluster sum of squares and between-cluster sum of squares, for various numbers of clusters. At around six clusters, increasing the number of clusters does not decrease this ratio as much; thus a division of the thalamus into six clusters was chosen for a deeper analysis. Resulting six clusters of thalamic voxels are shown on a **b)** coronal and **c)** axial slice; contours represent boundaries of thalamic nuclei in the Allen Mouse Brain Atlas (AMBA). Cluster-associated cortical structural covariance maps (average cortical structural covariance for all voxels within the thalamic cluster) are shown in **(d)** on sagittal, coronal, and axial slices; positions of each slice view of the brain are marked on the other two slice views. Further quantified for each cluster is **e)** the location of positive and negative cortical structural covariance, and **f)** the overlaps between thalamic clusters and AMBA-defined thalamic nuclei.

*Subdivisions of the thalamus.* The clusters of voxels that share similar cortical structural covariance profiles subdivide the thalamus into six regions of interest. **Figures 5b** and **5c** show these clusters along with the boundaries of AMBA-defined nuclei through coronal and axial slices respectively; for a more complete view, **Figures S5** (coronal) and **S6** (axial) show these data on closely-spaced slices that span the entire extent of the thalamus. All clusters contained between ∼10% and ∼20% of thalamic voxels, and most consist of a single major connected component (see **Supplementary Results**). Clusters can be labeled based on their location within the thalamus:

- cluster 1 (16183 voxels / 20.11% of the thalamus): ventral posterior,
- cluster 2 (8325 voxels / 10.35% of the thalamus): lateral,
- cluster 3 (16504 voxels / 20.51% of the thalamus): dorsal (epithalamus with intralaminar parts),
- cluster 4 (10109 voxels / 12.56% of the thalamus): ventral lateral,
- cluster 5 (12170 voxels / 15.13% of the thalamus): ventral anterior, and
- cluster 6 (17165 voxels / 21.33% of the thalamus): dorsal (mediodorsal nucleus with lateral parts).

*Spatial patterns of thalamocortical structural covariance.* The *k*-means cluster centers (average cortical structural covariance map for all thalamic voxels within the cluster) separate thalamocortical structural covariance into spatially distinct components. At six clusters—the optimal number of clusters as described above—cortical structural covariance components are very generally organized along the three cortical axes: anterior-posterior, medial-lateral, and superficial-deep. **Figure 5d** shows these average cortical structural covariance maps on sagittal, coronal, and axial slices that pass through the thalamic cluster, and **Figures S7** (sagittal), **S8** (coronal), and **S9** (axial) plot this same information on more slices that span the extent of the brain.

Each component is characterized by the presence of broad regions of both positive and negative cortical structural covariance (**Figure 5e**), and arise in pairs based on similarities in these patterns (**Supplementary Results**). In general, highest cortical structural covariance (positive and negative) was found in medial frontal areas (including the prelimbic, infralimbic, and orbital areas). Previously described depth-specific patterns separate across the six components; the separation of structural covariance by depth is particularly evident in medial parts of the frontal cortex, with some components showing high structural covariance in superficial layers, while other components show high structural covariance in deeper layers.

*Relationship between locations of thalamic clusters and cortical structural covariance maps.* We observed an interesting pattern in the spatial locations of thalamic clusters in that they generally reflected the organization of their cortical structural covariance components along the same axis (**Figure 5d**). For instance, clusters 1 and 5 are characterized by thalamic voxels in the ventral thalamus and high cortical structural covariance in deeper (ventral) layers. Conversely, clusters 3 and 6 consist of voxels in the dorsal thalamus and show high structural covariance in superficial (dorsal) layers. Clusters 2 and 4 consist of voxels in lateral parts of the thalamus, and are marked by high structural covariance in the lateral parts of the cortex.

*Thalamic subdivisions are weakly but significantly associated with anatomical definitions.* We computed the overlap of our structural covariance-based clusters of thalamic voxels with segmentations of the thalamus that are defined within the Allen Mouse Brain Atlas. For every pair of regions formed by pairing each AMBA structure with each of the six clusters, we independently computed three characteristic metrics that describe the overlap of the pair: the sensitivity (**Figure S10a**), positive predictive value (**Figure S10b**), and Dice coefficient (**Figure 5f**).

We found that overall, the overlaps as quantified by the Dice coefficient were not as high as those seen in classic studies on automated segmentations that seek to optimize labeling of well-defined regions on novel images (Chakravarty et al., 2013). Yet, in providing a new segmentation of the thalamus that defines regions based on a different information source (i.e. volume covariation patterns, as opposed to histology/cytoarchitecture), we find that these overlaps are statistically significant despite correcting for multiple comparisons via the conservative Bonferroni procedure that controls the family-wise error rate. For each cluster, we find statistically significant overlaps with multiple AMBA regions at all levels of the anatomical hierarchy, with only a 5% chance (under the assumptions of a permutation test) that at least one of these overlaps is a false discovery.

As a summary of these overlaps (**Figure 5f**), clusters 1, 2, and 4 overlap the sensory-motor cortex-related areas of the thalamus, while clusters 3 and 6 overlap the polymodal association cortex-related areas. The highest Dice coefficient was 0.434, corresponding to the overlap between cluster 3 and the AMBA-defined polymodal association cortex-related region. At a higher level of this tree, the greatest overlap was between cluster 6 and the medial group of the dorsal thalamus (MED), as quantified by a Dice coefficient of 0.363; this is mainly driven by the high sensitivity rather than positive predictive value (see **Supplementary Results** for a discussion on this). At this level, the structural covariance-based thalamic clusters overlap the following AMBA structures the best:

- cluster 1: Ventral group of the dorsal thalamus (Dice coefficient = 0.358),
- cluster 2: Lateral group of the dorsal thalamus (Dice coefficient = 0.275),
- cluster 3: Anterior group of the dorsal thalamus (Dice coefficient = 0.334),
- cluster 4: Geniculate group, dorsal thalamus (Dice coefficient = 0.337),
- cluster 5: Ventral group of the dorsal thalamus (Dice coefficient = 0.257), and
- cluster 6: Medial group of the dorsal thalamus (Dice coefficient = 0.363).

*Subdivisions of the thalamus at two clusters.* We also analyzed thalamocortical structural covariance at different numbers of clusters ranging from 2 to 40. At two clusters, cortical structural covariance is generally separated by depth, with high structural covariance present in deeper layers for cluster 1 and superficial layers for cluster 2 (**Figure S11**). High positive structural covariance was observed in frontal, cingulate, motor, and somatosensory areas (cluster 2) and in a similar set of regions that also included the prelimbic, infralimbic, and orbital areas (cluster 1) (**Figure S12**). Similar to the patterns observed when using six clusters, we found that 1) these clusters formed a pair based on the similarities between their positive and negative structural covariance profiles, and 2) the high structural covariance in the upper (dorsal) layers of the cortex were associated with the cluster of voxels in the dorsal thalamus, and vice versa (**Figure S11**). Compared to AMBA segmentations (**Figure S13**), these clusters best overlapped the two AMBA divisions of the thalamus at level 6 (**Figure S14**); cluster 2 best overlapped the polymodal association cortex-related region (Dice coefficient = 0.661), while cluster 1 best overlapped the sensory-motor cortex related-region (Dice coefficient = 0.465).

*Subdivisions of the thalamus at many clusters.* At six clusters, we observed significant but weak overlaps between our structural covariance-based thalamic clusters and the AMBA structures, which could be explained in part by a mismatch in the numbers (and consequently sizes) of the pairs of regions being compared. The Dice coefficient (overlap measure that we used) depends equally on both the sensitivity and positive predictive value, and when the sizes of the structures differ, at least one of these is penalized. Thus, we hypothesized that by increasing the number of clusters, there would be for each AMBA structure at least one thalamic cluster with a high overlap.

In the case of many clusters (up to 42), the thalamic subdivisions did not appear to approach the AMBA segmentations. For each AMBA structure, we computed the highest Dice coefficient across all cluster numbers and cluster assignments, and found that the majority (∼80%) of these values were below 0.4 (**Figure S15**). The highest overlaps observed (for AMBA structures at hierarchy level 7 or above) involve the epithalamus (which includes the medial and lateral habenula, maximum Dice coefficient for the epithalamus = 0.805) and the mediodorsal nucleus (maximum Dice coefficient = 0.576). For the epithalamus, the highest Dice coefficient was found at 28 clusters, although similar large overlap values were present from 20 to 40 clusters. In the case of the mediodorsal nucleus, the highest Dice coefficient occured at 22 clusters, corresponding to a segmentation that forms by splitting the aforementioned cluster 6 that is obtained when using six clusters.

Structural covariance-based thalamic subdivisions and associated cortical structural covariance maps for 2 to 42 clusters are provided as MINC and NIFTI files within the **Supplementary Data**.

### 3.3. Fiber tracts associated with thalamocortical structural covariance

We further examined whether the six clusters of thalamic voxels and associated cortical structural covariance maps reflected cluster-specific thalamocortical structural connectivity. For each cluster, we computed the overlap between thalamic voxels and injection sites from the Allen Institute’s anterograde viral tracing data, and thus found a series of “experiments” of tracer projections to the cortex that emanate from each thalamic cluster. A total of 171 such experiments were found across all clusters. Thalamic clusters 1 to 6 overlapped 28, 18, 41, 14, 15, and 55 injection sites respectively, providing sufficient coverage of thalamocortical structural connectivity.

For each cluster, we merged all associated tracer projection images by taking the maximum projection density at each voxel, thereby creating a single image that describes cluster-specific thalamocortical connectivity (**Figure 6a**). We found that cortical areas that expressed the tracer also tended to have high structural covariance, indicating that cortical structural covariance associated with the thalamic clusters is sensitive to connectivity. We note that this cortical structural covariance is not specific to structurally connected areas however; high structural covariance is also observed in cortical regions unconnected to the thalamic clusters.

**Figure 6:**
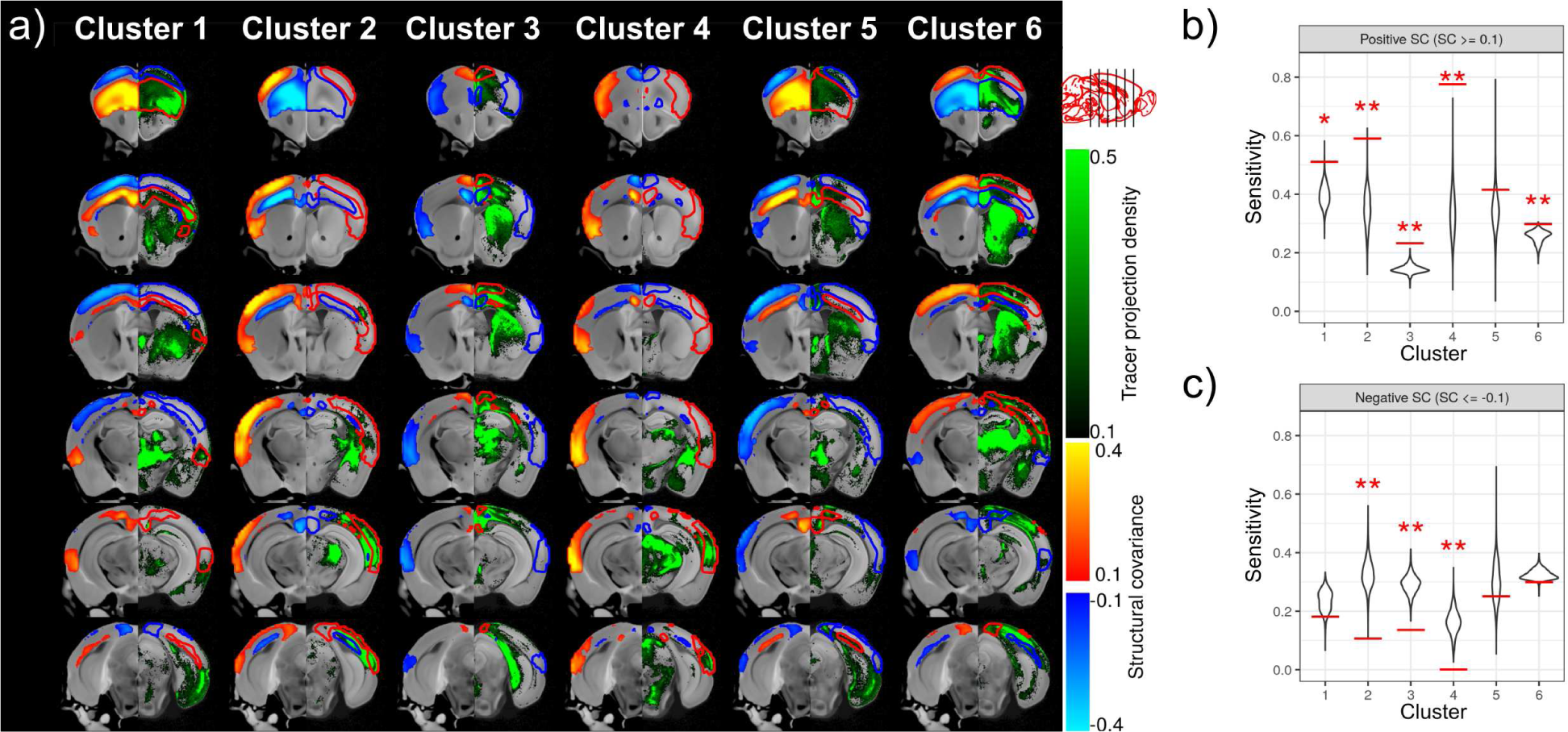
Association between thalamocortical structural connectivity and structural covariance. **a)** For each cluster as a column of coronal slices, cortical structural covariance (left hemisphere) is compared to a maximum-intensity projection of all tracer projection density images that result from injections within the cluster (right hemisphere). Contours constructed from thresholding the structural covariance data are also provided as a visual guide on the right. The overlaps between thalamocortical structural connectivity and structural covariance are further quantified via permutation tests for **b)** positive and **c)** negative structural covariance. For each cluster, the overlap between regions of high absolute cortical structural covariance (positive and negative) and cortical regions innervated by tracer projections that arise from the thalamic cluster are assessed as the sensitivity (red lines). These are overlaid on permutation distributions, formed by spatially permuting cluster definitions and recomputing the sensitivity statistic; *p*-values are computed as twice the proportion of permuted statistics that are more extreme than the observed statistic. Here, sensitivity is assessed by considering the viral tracer images as ground truth, i.e. sensitivity is quantified as the proportion of cortical voxels connected to thalamic clusters that are also overlapped by regions of high structural covariance. Tracer projection density images are thresholded at 0.1; cortical structural covariance maps are thresholded at 0.1 (positive) or −0.1 (negative). Statistically significant relationships (after adjusting *p*-values for multiple comparisons via the FDR method) are annotated: * *p* < 0.05, ** *p <* 0.01.

Using permutation tests, we found that for all clusters except cluster 5, the sensitivity of high positive structural covariance to structural connectivity was statistically significant (**Figure 6b**). Conversely, for negative structural covariance, sensitivity was far lower than could be explained by randomness for clusters 2 to 4, indicating that negative structural covariance tends to be preferentially located in unconnected regions (**Figure 6c**). FDR-corrected *p*-values are provided in **Table 2** below.

**Table 2:** *p*-values associated with a permutation test of the sensitivity of structural covariance to structural connectivity. *p*-values are provided for both positive and negative structural covariance, and are corrected for multiple comparisons using the FDR method.

| Structural covariance sign | Cluster 1 | Cluster 2 | Cluster 3 | Cluster 4 | Cluster 5 | Cluster 6 |
| --- | --- | --- | --- | --- | --- | --- |
| Positive | 0.0106 | 0.00240 | 0.00120 | 0.00120 | 0.216 | 0.00900 |
| Negative | 0.150 | 0.00160 | 0.00120 | 0.00120 | 0.480 | 0.0666 |

## 4. Discussion

### 4.1. Subdivisions of the thalamus

Building on previous studies by others that subdivide the thalamus based on structural (Behrens et al., 2003; Johansen-Berg et al., 2005; Traynor et al., 2010; O’Muircheartaigh et al., 2011; Lambert et al., 2017; Middlebrooks et al., 2018) and functional (Zhang et al., 2008; Toulmin et al., 2015; Yuan et al., 2016; Ji et al., 2016; Long et al., 2021) connectivity, and examine subdivisions in other structures based on structural covariance (Cohen et al., 2008; Kelly et al., 2012; Plachti et al., 2019), we compute cortical structural covariance maps for each voxel in the thalamus as a seed, and find that these voxels cluster into six components. Grouping the 80456 cortical structural covariance maps into six clusters provides insights into the spatial organization of thalamocortical structural covariance. We found that the patterns of cortical structural covariance were similar to the locations of thalamic clusters along each of the three standard axes. For clusters that showed high cortical structural covariance in superficial (dorsal) layers of the cortex, the associated set of thalamic voxels were in the dorsal part, and vice versa. Similar patterns that indicated an anterior-posterior and medial-lateral organization were also seen. While the exact mechanisms behind this organization are unclear, the cluster-specific association between thalamocortical structural covariance and structural connectivity suggests that wiring may play a role. For example, these patterns would be seen if 1) wiring costs require that distances between thalamic clusters with areas of high cortical structural covariance be minimized, and 2) there exists variation in the numbers and sizes of neurons forming these tracts.

The dorsal-ventral split appeared to be the most important, since a clustering of the data into two clusters subdivided the thalamus along this axis. Here too, cortical structural covariance is organized into two components that stratify along this axis. It is interesting that the AMBA subdivision of the thalamus into polymodal association cortex-related areas and sensory-motor cortex-related areas also subdivide the thalamus along the dorsal-ventral axis. The structural covariance-based cluster that overlaps the association cortex-related nuclei is associated with high structural covariance in the superficial layers of the cortex, while the structural covariance-based cluster that overlaps the sensory-motor cortex-related nuclei is associated with high cortical structural covariance in deeper layers; these could reflect the fact that upper layers integrate information between different areas through corticocortical projections, while lower layers relay information downstream via projections to subcortical structures. We note that this split is also consistent with the cortical aspect of the core-matrix framework which divides thalamic projections into a “core” group of neurons that terminate in the lower layers and a “matrix” group of neurons that terminate in the upper layers. The former group is thought to display a sharp response to stimuli and relay information to precise areas, while the latter group is thought to facilitate the integration of different functions and support consciousness (Jones, 1998). Connecting thalamocortical structural covariance to projection cell types would be helpful in further understanding the aforementioned organizational principles. Many of the Allen Institute projection experiments were carried out in Cre-lines and thus label specific projection cell types (Harris et al., 2019); these may help identify which cell types contribute to thalamocortical structural covariance and explain why it varies across the cortical depth.

It is worth noting that when comparing our six subdivisions of the thalamus to AMBA segmentations, we found generally weak overlaps (as quantified by the Dice coefficient), with the epithalamus and mediodorsal nucleus being exceptions. We hypothesize that this is due to a combination of two factors. First, structural covariance-based segmentations might reflect an alternative subdivision of the thalamus that do not correlate with traditional cytoarchitecturally-defined nuclei, and may instead better reflect gene expression patterns (Yee et al., 2018) which do not necessarily obey traditionally defined boundaries. For example, the core-matrix framework proposes that matrix-related D28K Calbindin expressing cells are distributed in the thalamus in a non-specific manner (Jones, 1998), and recent RNA sequencing of thalamocortical projections suggests that there are three major transcription profiles that are seen across all thalamocortical functional systems (Phillips et al., 2019). Second, it is difficult to resolve individual nuclei with MRI even with optimized sequences (Magnotta et al., 2000). Some thalamic subdivisions are visible in the *T*_2_-weighted images used here, but the mapping of fine nuclei might require an approach that combines multimodal imaging data. This explains the case of the epithalamus and mediodorsal nucleus, which are better outlined by structural covariance-based clusters; these two structures are anatomically separated from rest by intralaminar nuclei and they themselves via the fasciculus retroflexus, and can be discerned through *T*_2_-weighted MRI as seen in **Figure S1a**.

Within the few studies that have subdivided brain structures using structural covariance, both convergence and divergence of structural covariance-based segmentations with other types of connectivity-based segmentations have been reported. Specifically, Kelly et al. (2012) report that structural covariance-based segmentations of the human insula are similar to ones based on functional connectivity, while Plachti et al. (2019) observe a weaker concordance in the human hippocampus. While we do not examine the concordance between our clustering of thalamic voxels with rs-fMRI or DTI-based segmentations, our results are closer to the latter case (the AMBA segmentations of the thalamus which we compare our results to are delineated in part by structural connectivity, as assessed by neuronal tracers). In the hippocampus, Plachti et al. (2019) observe a medial-lateral organization of structural covariance-based subdivisions that is thought to map to cytoarchitectural differences between CA subfields and the subiculum; this is in contrast to rs-fMRI and meta-analytic connectivity modeling-based subdivisions that are dominantly organized along an anterior-posterior axis. Similar to the hippocampus in which these structural covariance modes of variation along different axes are attributed to different functional and microstructural properties (Masouleh et al., 2020), the organization of our thalamic segmentations along each axis may be related to separate factors which are yet to be determined.

### 4.2. Negative structural covariance

Another observation in our study worth noting is the presence of high negative structural covariance, preferentially located in areas of the cortex unconnected to the thalamus, which could in part be explained by data normalization. Interestingly, components could be paired based on similarities in their cortical structural covariance maps, with each positive structural covariance profile being paired with a negative one. Such a pairing could presumably arise from the regression of parent structure (cortex and thalamus) volumes from the Jacobian determinants, in a way that is analogous to the anticorrelations that are seen in fMRI analysis after global signal regression (Fox et al., 2009; Weissenbacher et al., 2009). Negative structural covariance might also arise from functional competition between regions however, which in turn could be amplified in magnitude through the data normalization. For example, anticorrelations in volume have been observed between the hippocampus and striatum (structures which are associated with competing strategies used in learning to navigate) (Bohbot et al., 2007). It is unclear how functional competition would explain the variation in structural covariance along layers however, since differences in learning strategies would be associated with differences in areas rather than layers. While we consider positive and negative structural covariance on an equal footing in clustering the data, more work must be done to understand the effects of data processing on correlational networks (Liang et al., 2012; Zalesky et al., 2012), and the extent to which these anticorrelations in volumes reflect biology.

### 4.3. Growth or connectivity?

What might cause these thalamocortical structural covariance networks to form? One explanation is the “wire together, fire together, grow together” hypothesis: regions that are connected tend to coactivate and undergo plastic changes in a coordinated manner (Evans, 2013); variation in the extent of this coactivation across the population would then explain the formation of structural covariance. Consistent with this are our observations that thalamocortical structural covariance is sensitive to structural connectivity and is seen in areas of the cortex that are functionally linked to the chosen seeds. Structural covariance is not predictive of connectivity however, and one explanation for this could be that volume covariance reflects functional connectivity acting over broader networks, while the viral tracing data we have used maps the paths of tracers that do not cross synapses (Oh et al., 2014) and therefore describes a much more limited type of brain network. Although the broader patterns of structural covariance could be explained by connectivity and subsequent coactivation across multiple synapses, previous studies have shown that computationally-estimated connectivity across multiple synapses (using the same viral tracing data) does not do much better in explaining structural volume covariance (Yee et al., 2018).

Emerging evidence suggests that structural covariance might also form from the coordinated growth of structures that undergo similar developmental processes (Alexander-Bloch et al., 2013b), perhaps due to the expression of common sets of genes (Romero-Garcia et al., 2018; Yee et al., 2018). The observation of a spatial correlation between connectivity and structural covariance describes an association rather than a cause, and is consistent with this latter hypothesis; regions that are connected tend to develop together (Raznahan et al., 2011) and express similar gene expression profiles (French & Pavlidis, 2011; Richiardi et al., 2015). Thus, structural covariance might develop in conjunction with (or perhaps even before) structural and functional networks, for example as a byproduct of variation in the numbers and sizes of cells in precursor structures, or in the neurons that connect them, as measured across the population. We note that the coordinated neurodevelopment hypothesis does not preclude connectivity and activity, since cortical patterning (and presumably volumes) are tightly coupled to thalamic development (López-Bendito & Molnár, 2003). Specifically, cortical arealization is thought to partly depend on thalamic inputs (Rakic, 1988; Krubitzer & Huffman, 2000), and the coordinated neurodevelopment hypothesis suggests that it is during these early critical periods of neurodevelopment that thalamocortical structural covariance forms.

We further note that in this study, thalamocortical structural covariance was determined in a group of inbred mice of the same C57Bl/6 strain (though notably, from two different substrains, B6J and B6N, see **Table 1**). Mice were raised in standard cage conditions, and were within a narrow age range (postnatal day 56-84 corresponding to early adulthood) during which brain anatomy is not expected to change much. Despite very little genetic, environmental, and age-related variation in this population, we observed structural covariance, leaving room for variation at the transcriptomic level (e.g. in response to differences in maternal care or the intrauterine environment) and stochastic variation as further explanations. In contrast, both male and female mice were included, and this might be a source of variation in anatomy. Here, we chose to pool both sexes together in order to examine structural covariance reflective of the entire population. Sex-related differences in structural covariance have been reported by others before (Persson et al., 2014), but are outside the scope of our study. In terms of environmental variation driving structural covariance, a recent study demonstrated that mice raised in an enriched environment showed stronger structural covariance networks, as characterized by generally higher correlations between the volumes of pairs of regions (Bogado Lopes et al., 2023). A similar study with a focus on thalamocortical structural covariance involving outbred mice and/or those raised in enriched environments would be welcome; we hypothesize that in such studies, more components (than the six seen here) would emerge. Furthermore, given the extensive knowledge of spatial gene expression in the developing mouse brain (Thompson et al., 2014) and the timing of neurodevelopmental processes in mammals (Andersen, 2003), a longitudinal study that examines timecourses of thalamocortical structural covariance would shed light on the role of coordinated growth and experience-dependent plasticity as causal mechanisms.

### 4.4. Methods and limitations

We use volume as a structural property from which covariation patterns are computed. Compared to cortical thickness— a structural measure commonly used in studies on human structural covariance—volumes are generally easier to compute in the mouse (due to the relative ease of registering lissencephalic brains and generating brainwide Jacobian determinant maps via deformation-based morphometry), and are defined for both the cortex and thalamus. In the cortex, volume can be seen as the product of cortical thickness and surface area; thickness and area are orthogonal measures that are shaped by distinct genetic mechanisms, differ in heritability, and are influenced by the environment to different extents (Sanabria-Diaz et al., 2010; Winkler et al., 2010; Joshi et al., 2011; Eyler et al., 2012; Jha et al., 2018; Strike et al., 2019). Correlational networks constructed from these different measures demonstrate little overlap in their edges and display different topologies (Sanabria-Diaz et al., 2010; Yang et al., 2016). Therefore caution must be used when interpreting these volume-related results in relation to other studies involving cortical thickness. Although it is possible to use separate structural measures for the cortex and thalamus (e.g. cortical thickness and thalamus volume), our observation on the variance of cortical volume covariance along the depth-axis suggests that cortical thickness (which integrates volumes along streamlines over all layers) would remove this important signal. Due to this reason, prior attempts at subdividing the thalamus using cortical regions of interest that spanned all layers were unsuccessful. Finally, due to the lack of clear areal boundaries in lissencephalic brains, using surface area was not feasible.

Estimating volumes relies on a registration process that works by spatially averaging structures based on contrast. In our case, a fast spin echo sequence provided *T*_2_-weighted contrast (Spencer Noakes et al., 2017). Consequently, volume estimates will not be accurate for regions in which there is little to no such contrast, and the subsequent estimates of covariance will be flawed. If *T*_2_-weighted contrast reflects cell density and myelin content (Boretius et al., 2009), then volume changes in regions that share a boundary with others of similar cell density and myelination may not be accurately captured. A similar study using images of different contrasts would be beneficial in understanding the role of the MR sequence in the estimation of structural covariance networks.

As mentioned previously, our study is further limited by the use of mice that display little genetic and environmental variation, thus limiting the signal from which structural covariance is estimated. Furthermore, volumes are processed by regressing out cortex/thalamus volumes from Jacobian determinants. While this overall scaling due to parent structure size is unwanted, this regression is analogous to the global signal regression problem that arises in fMRI analyses in which the signal-to-noise ratio is diminished, and is likely part of the reason why large negative values of structural covariance are observed.

### 4.5. Future directions

Beyond replicating such an analysis using outbred mice and in longitudinal datasets, similar studies using human imaging data could provide components that are useful in the clinic. Although structural covariance is a group-wise measure, individual “integrity scores” may be computed by projecting patient cortical measures onto these standard components. Of particular interest is the relationship between cortical structural covariance and depth. Given the additional complexity of the human cortex relative to the mouse, examining this depth dependence in humans while also understanding the cellular correlates of this in the mouse would be of translational value. Interrogating these results using spatial gene expression data could help identify the sets of genes, cell types, and developmental mechanisms that are involved in shaping thalamocortical structural covariance and its relationship to cortical layers; these genes and cell types can then be knocked out as part of a further validation study.

Finally, we note that we examined thalamocortical structural covariance using a clustering approach. An alternative and equally valid approach would be to characterize gradients (Masouleh et al., 2020; Bernhardt et al., 2022) of structural covariance within the thalamus. Code and data are available for other researchers if they wish to re-analyze thalamocortical structural covariance using such other frameworks.

### 4.6. Conclusion

In this study, we confirm previous findings and describe several novel findings. First, we demonstrate that structural covariance between the local volumes within the thalamus and cortex reflects known functional connections. Consistent with previous studies that connect structural covariance with structural connectivity, we report that cortical structural covariance to thalamic seeds are sensitive to structural connectivity, but are not predictive of it, and can be found in broader sets of regions. Then, we newly show that cortical structural covariance maps vary as a function of cortical depth. This relationship with cortical depth has important implications on the use of measures that integrate along cortical streamlines, suggesting that measures such as cortical thickness (computed entirely between the pial and white matter surfaces) will average away the cortical structural covariance signal, and that measures more localized along the depth axis are better suited to examine cortical structural covariance. Finally, we observed sharp transitions in cortical structural covariance maps when thalamic seed location was varied, and demonstrate that the thalamus is best clustered into six groups of voxels. While these clusters weakly associate with certain thalamic nuclei, our structural covariance-based segmentations of the thalamus appear to reflect a more general organizational principle of morphometric covariation in that both thalamic clusters and their cortical structural covariance maps organize along the dorsal-ventral, anterior-posterior, and medial-lateral axes.

For future studies on structural covariance in mice, we provide these structural covariance-based segmentations and their associated cortical structural covariance maps (for total cluster numbers ranging from 2 to 42), all of which are defined over the Allen Institute’s commonly used CCFv3 template. By computing volumes under each thalamic cluster and projecting cortical volumes onto cortical structural covariance maps, the integrity of these canonical components of coordinated cortical and thalamic variation can be assessed in mouse models as biomarkers of disease and treatment.

## Acknowledgments

This work was supported by funding from the Canadian Institutes of Health Research, Brain Canada, and the Ontario Brain Institute. We thank Benjamin Darwin for discussions and guidance on the image registration pipeline used in this study. We additionally thank the Allen Institute for Brain Science for providing the connectivity (connectivity.brain-map.org) and reference atlas data (atlas.brain-map.org) used here; details on these datasets are contained within their primary publications cited in the Methods.

## Supplementary Data

Attached as Supplementary Data:

- Movies related to **Figures 4** and **S4**
- Structural covariance-based segmentations of the thalamus (2-42 clusters)
- Aligned templates (CCFv3 and MRI) and masks

## Supplementary Results

### Subdivisions of the thalamus

*Most clusters consist of a single major connected component.* All clusters except for cluster 6 consist of a single major connected component (defined as containing at least 90% of all voxels assigned to the cluster), with the remaining few voxels generally being at the boundary of the thalamus. In the case of cluster 6, voxels within the cluster form two major connected components that outline 1) the lateral part of the dorsal thalamus (65.30% voxels in cluster) and 2) the mediodorsal nucleus (34.69% of voxels in cluster). Visually, the overlap between the mediodorsal nucleus and this second connected component of cluster 6 was very clear. Of the remaining voxels not part of the major connected components of any cluster, the largest component formed was only 1.91% of all cluster-assigned voxels (cluster 4). While these voxels are retained in subsequent analyses comparing clusters to AMBA segmentations and structural connectivity, their limited numbers mean that they contribute little to the quantification and interpretation of these clusters.

*Components can be paired based on cortical structural covariance similarities.* An intriguing pattern is that components can be paired based on similarities in their cortical structural covariance profiles (**Figure 5e**). In the case of positive structural covariance:

- components 1 and 5 show high structural covariance in the deeper layers of the frontal cortex (including the orbital, prelimbic, infralimbic, anterior cingulate, motor, and agranular insular areas) along with weaker structural covariance in the retrosplenial and visual areas,
- components 2 and 4 show high structural covariance in the superficial layers of the parietal and temporal areas (including the somatosensory, auditory, gustatory, visceral, temporal association, perirhinal, and ectorhinal areas), and
- components 3 and 6 show high structural covariance in the superficial layers of medial frontal areas (in areas similar to components 1 and 6 but excluding the infralimbic area).

In the case of negative structural covariance, components can also be paired, though the pairs are different. The differences in pairing observed here (as compared to the positive structural covariance case) are responsible for the separation of cortical structural covariance into six clusters; if the pairs were the same, the optimal number of clusters would be reduced to three. For negative structural covariance:

- components 1 and 4 show high negative structural covariance in the superficial layers of the medial cortex (including the motor, somatosensory, anterior cingulate, and retrosplenial areas),
- components 2 and 6 show high negative structural covariance in the deeper layers of the medial cortex (the same regions as components 1 and 5 along with the addition of the frontal, orbital, prelimbic, and infralimbic areas), and
- components 3 and 5 show high negative structural covariance in parietal and temporal areas (including the somatosensory, auditory, gustatory, visceral, temporal association, perirhinal, and ectorhinal areas).

Furthermore, each component could also be matched with another by pairing positive and negative structural covariance profiles. Specifically, components showing similar structural covariance patterns of opposing sign are:

- components 1 and 6,
- components 2 and 5, and
- components 3 and 4.

For each of these pairs, the positive structural covariance profile of one component is similar to its pair’s negative structural covariance profile and vice versa. It is interesting that across these three sets of pairings, there is no pair of components that is the same. For any one component, the spatial patterns of positive and negative structural covariance are different.

*Optimizing sensitivity vs. positive predictive value.* Given that we compare our thalamic clusters to AMBA segmentations that are at various levels of a hierarchy of structures and thus range in size, sensitivity values were generally higher for regions at the largest level of the anatomy tree (since these regions were small enough to be entirely covered by the structural covariance-based clusters, **Figure S10a**). For cluster 1 overlapping the ventral posterolateral nucleus of the thalamus, parvicellular part (VPLpc), and for cluster 4 overlapping the intergeniculate leaflet of the lateral geniculate complex (IGL), the subgeniculate nucleus (SubG), and the intermediate geniculate nucleus (IntG), the sensitivity reached unity. For similar reasons, the positive predictive values were generally highest for structures closest to the root (thalamus) (**Figure S10b**). Highest positive predictive values of the thalamic clusters were for clusters 3 and 6 predicting the polymodal association cortex-related region as defined at level 6 of the AMBA hierarchy (0.849 and 0.759 respectively).

## Supplementary Figures

**Figure S1:**
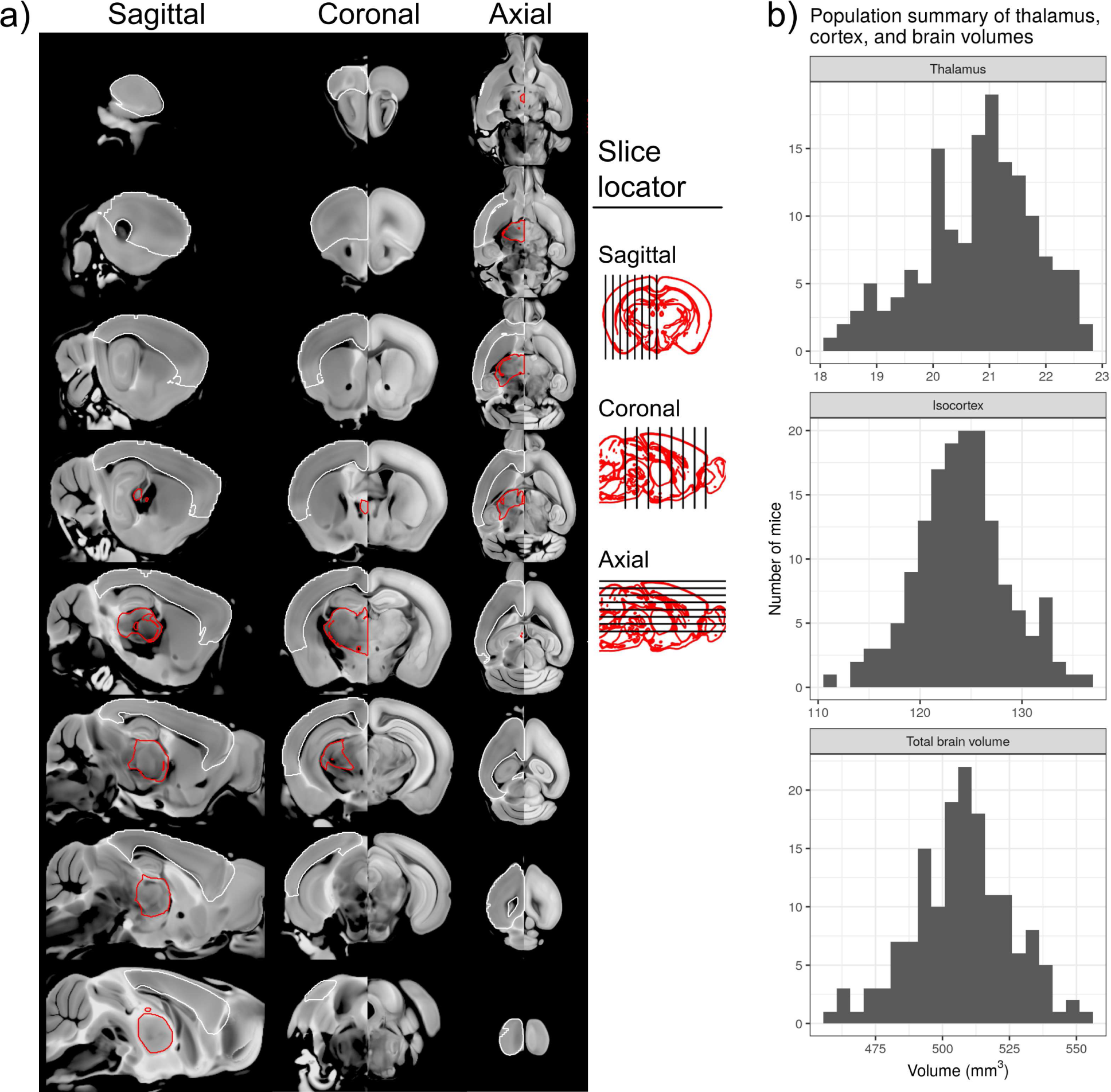
A visualization of the MRI to CCFv3 registration quality, the masks within which thalamocortical structural covariance is computed, and volumes of relevant structures. In **a)** sagittal, coronal, and axial slices through the mouse brain are presented, with both MRI average (left hemisphere) and CCFv3 template (right hemisphere) displayed together to visualize MRI to CCFv3 registration quality. In computing structural covariance, only the left hemisphere is considered. Shown are boundaries of the thalamus mask (red) under which each 50 *µ*m-resolution voxel was used as a seed, and the 200 *µ*m mask (white; resampled at 50 *µ*m for visualization) under which cortical structural covariance was computed for each thalamic voxel as a seed. Both masks were first intersected with another mask under which gene expression (coronal dataset from the Allen Brain Atlas) is defined, thus making future studies that involve linking cortical structural covariance to gene expression data easier to perform. In **b)**, the distribution of volumes of the thalamus, isocortex, and whole brain (both hemispheres) over all 154 subjects are shown.

**Figure S2:**
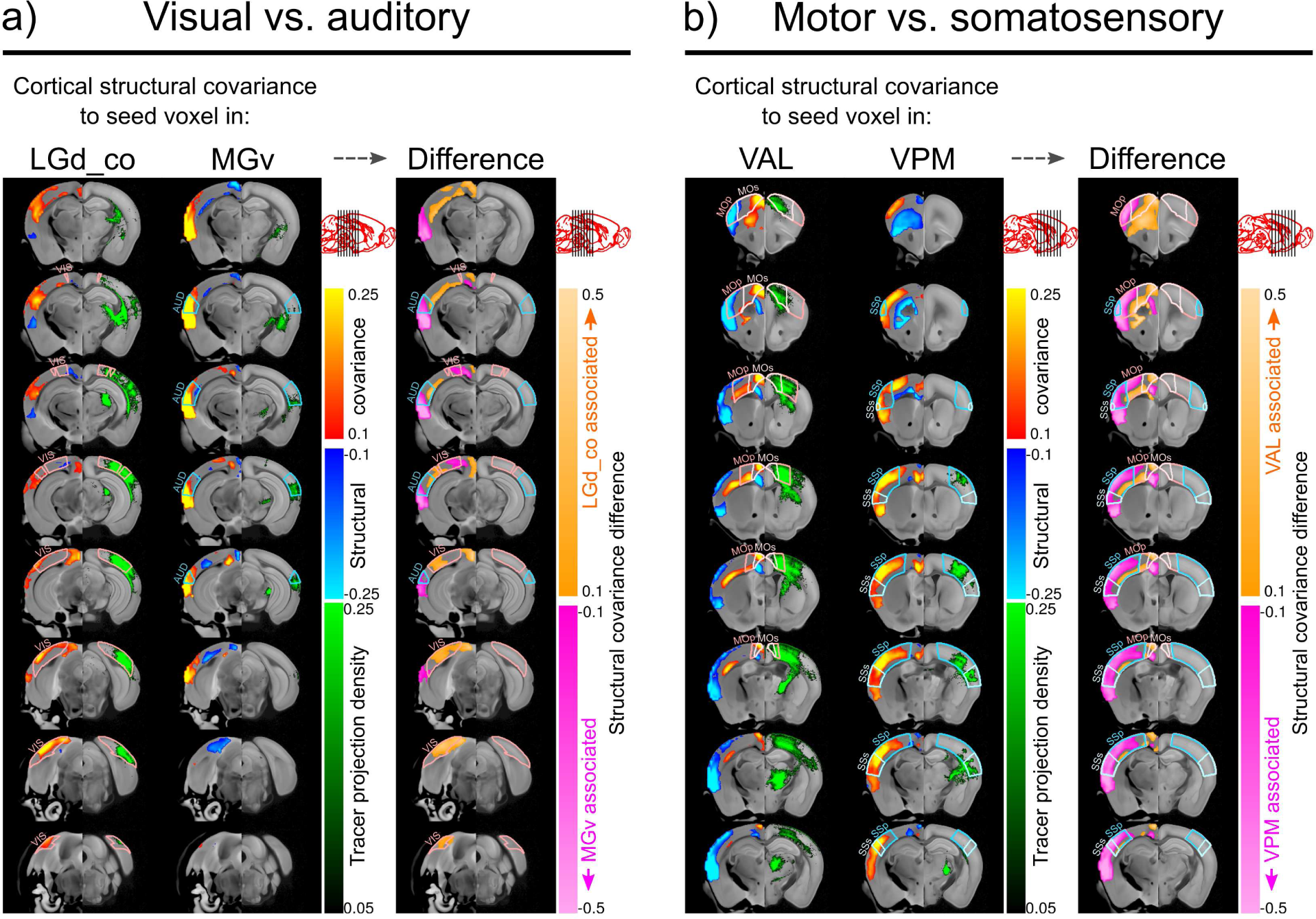
Seed-based cortical structural covariance maps corresponding to four functionally-relevant seeds (same data as **Figure 2**), plotted over a 3D volume. In **a)** LGd- and MGv-related structural covariance representing visual and auditory systems respectively are compared, while in **b)** VAL- and VPM-related structural covariance representing motor and somatosensory systems respectively are compared. Columns of coronal slices show cortical structural covariance (computed within a left-hemisphere cortical mask and plotted over the MRI template) for a seed voxel representative of each system. For comparison, data from the Allen Institute are plotted in the opposite (right) hemisphere: the CCFv3 template is plotted as the background, along with an overlay of a viral tracer that was injected in the structure of interest. Regions of interest based on the Allen Mouse Brain Atlas segmentations are outlined on both hemispheres. Difference in cortical structural covariance between these two maps highlight the spatial specificity of thalamocortical structural covariance in contrasting functional systems.

**Figure S3:**
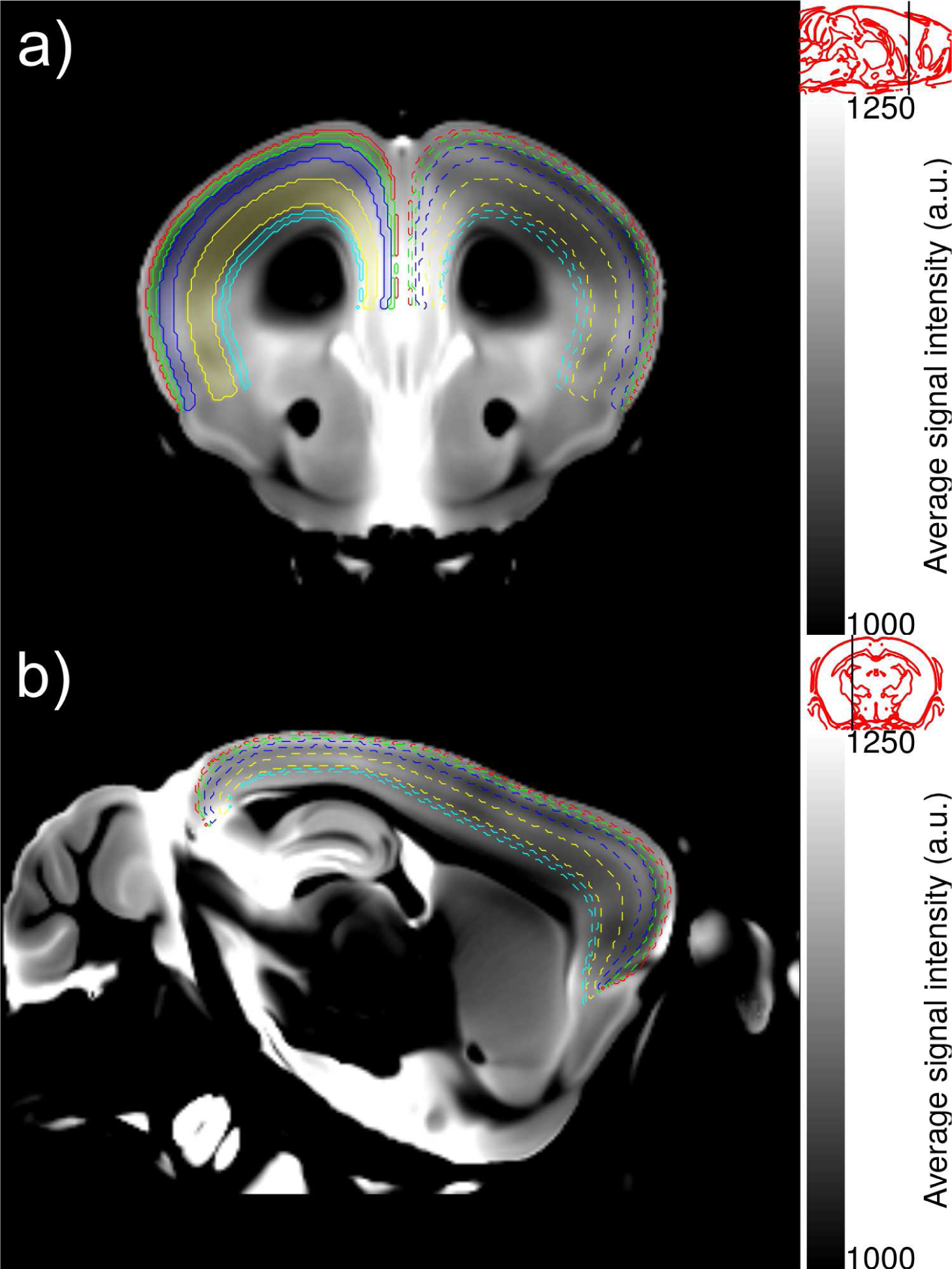
Layer-related contrast in an average *T*_2_-weighted image from which depth-dependence of cortical structural covariance can be analyzed. Highlighted on a **a)** coronal and **b)** sagittal slice of the MRI study template are the five major depths at which high structural covariance is observed (see **Figure 3d**). The MR (postprocessed) image intensities are scaled to increase contrast and depict the structural information from which Jacobian determinants (local volumes) are computed.

**Figure S4:**
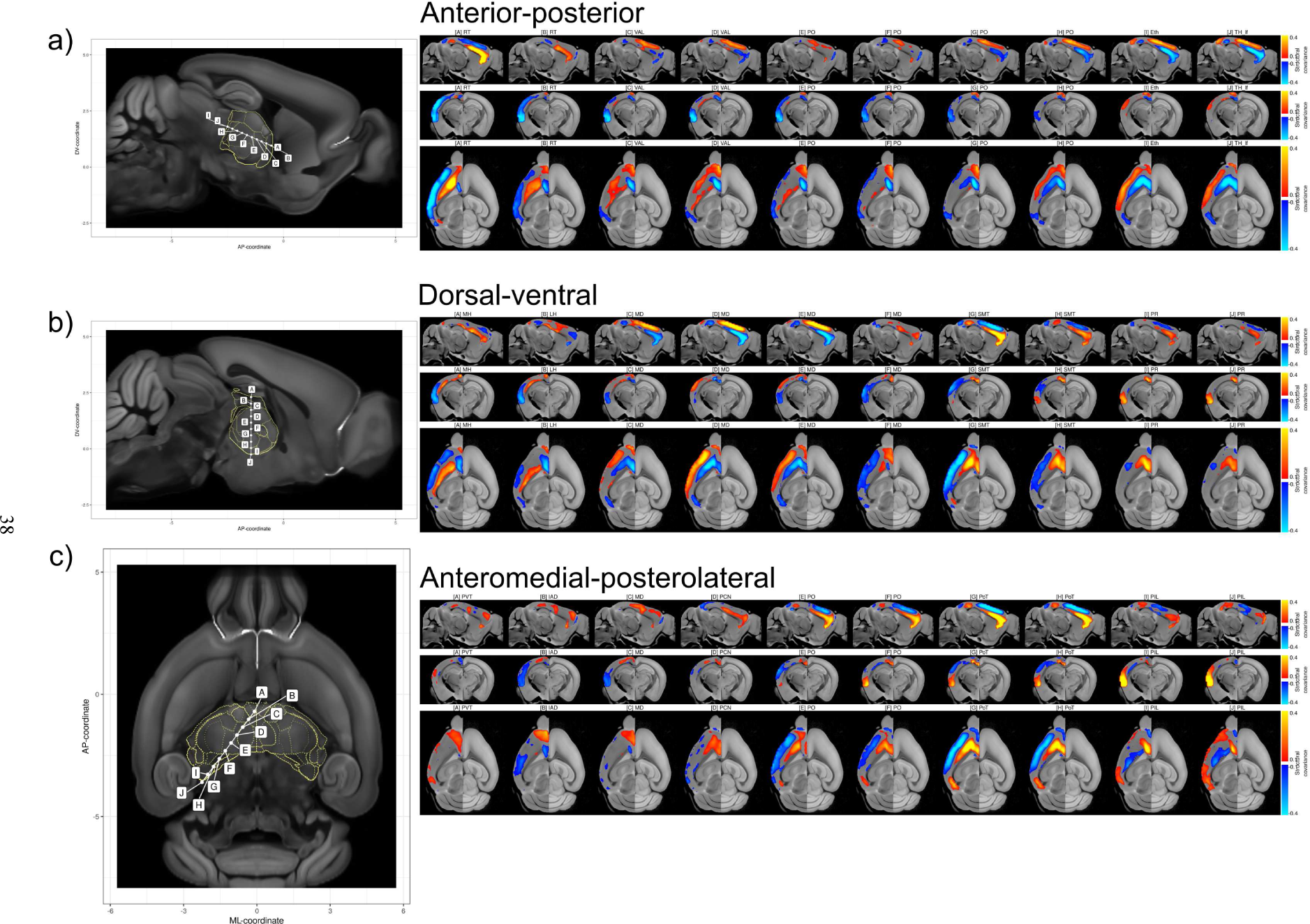
Cortical structural covariance for various equally-spaced seed points in the thalamus that are arranged along **a)** an anterior-posterior axis, **b)** a dorsal-ventral axis, and **c)** an anteromedial-posterolateral axis. Each annotated seed point (left panel) corresponds to a cortical structural covariance map that is plotted on sagittal, coronal, and axial slices (right panel). Labels at the top describe the thalamic nucleus (denoted by its Allen Mouse Brain Atlas acronym) that the seed point lies within.

**Figure S5:**
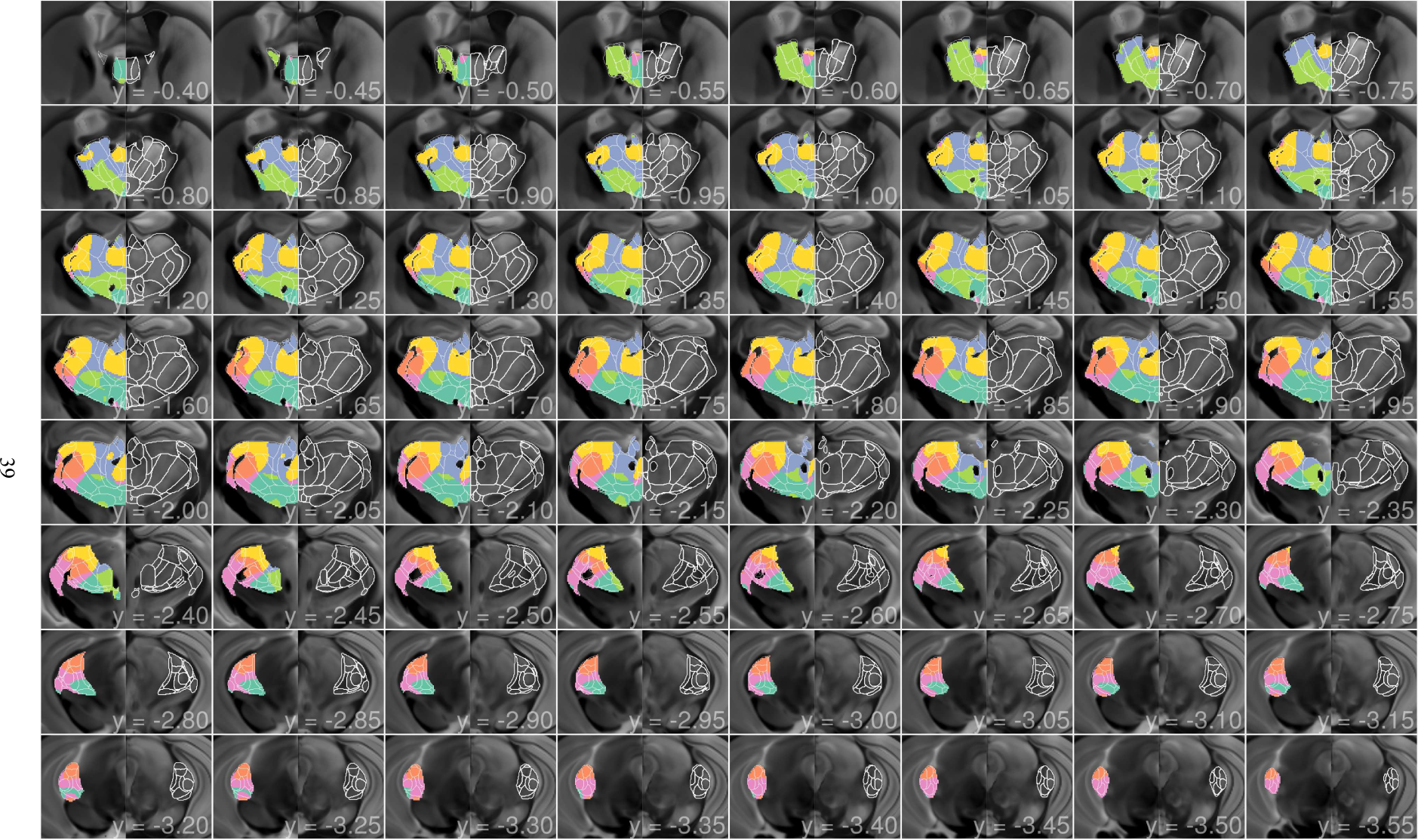
A coronal view of cortical structural covariance-based subdivisions of the thalamus into six clusters. A panel of coronal slices show the thalamus from the anterior to posterior end. Within each slice, covariance-based subdivisions are shown in the left hemisphere on the study MRI average as the background, and contralateral to this, thalamic anatomy is visualized via the CCFv3 template in the right hemisphere. Allen Mouse Brain Atlas segmentations of thalamic nuclei are represented via boundary contours in both hemispheres.

**Figure S6:**
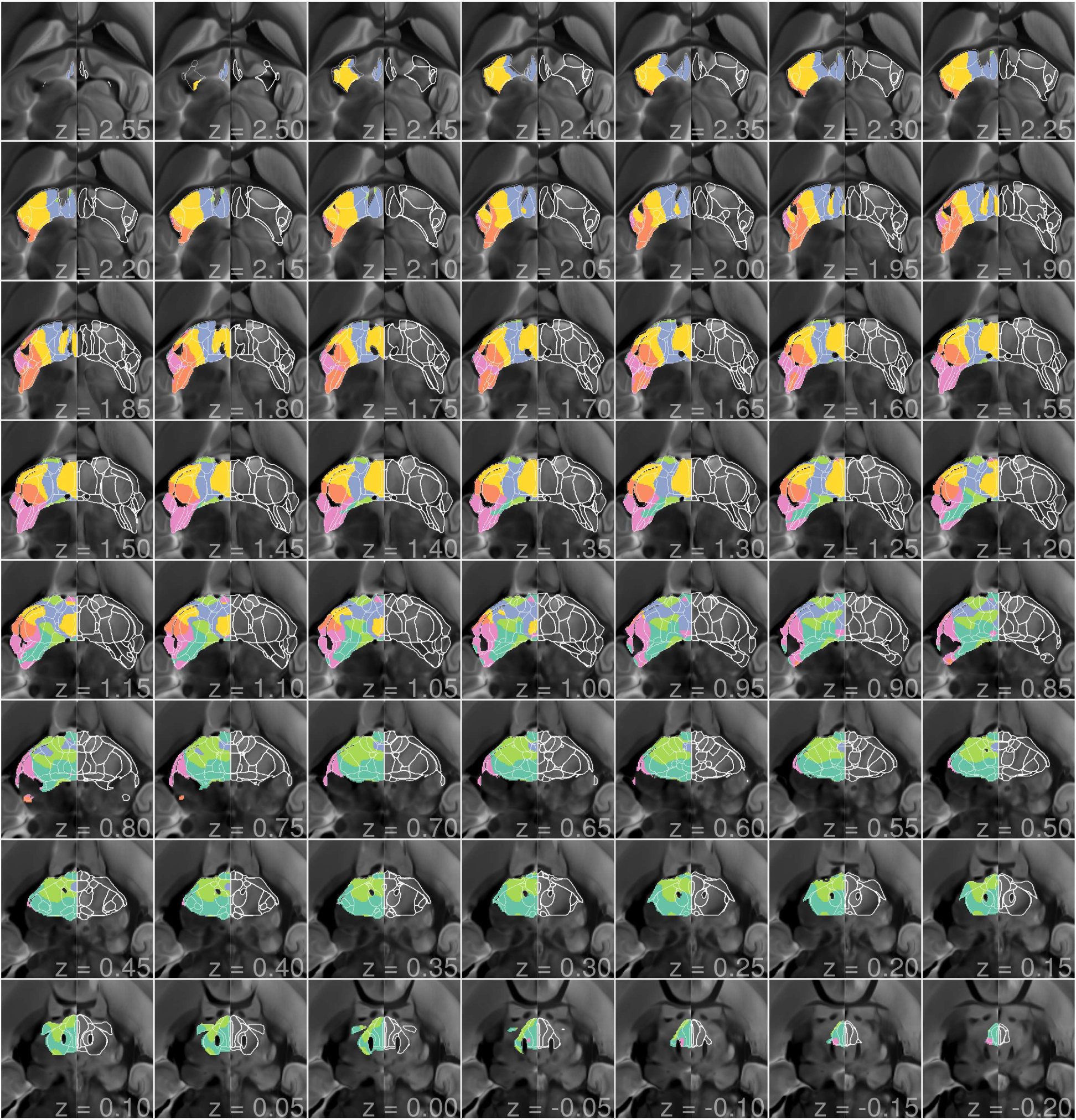
An axial view of cortical structural covariance-based subdivisions of the thalamus into six clusters. A panel of axial slices show the thalamus from the dorsal to ventral end. Within each slice, covariance-based subdivisions are shown in the left hemisphere on the study MRI average as the background, and contralateral to this, thalamic anatomy is visualized via the CCFv3 template in the right hemisphere. Allen Mouse Brain Atlas segmentations of thalamic nuclei are represented via boundary contours in both hemispheres.

**Figure S7:**
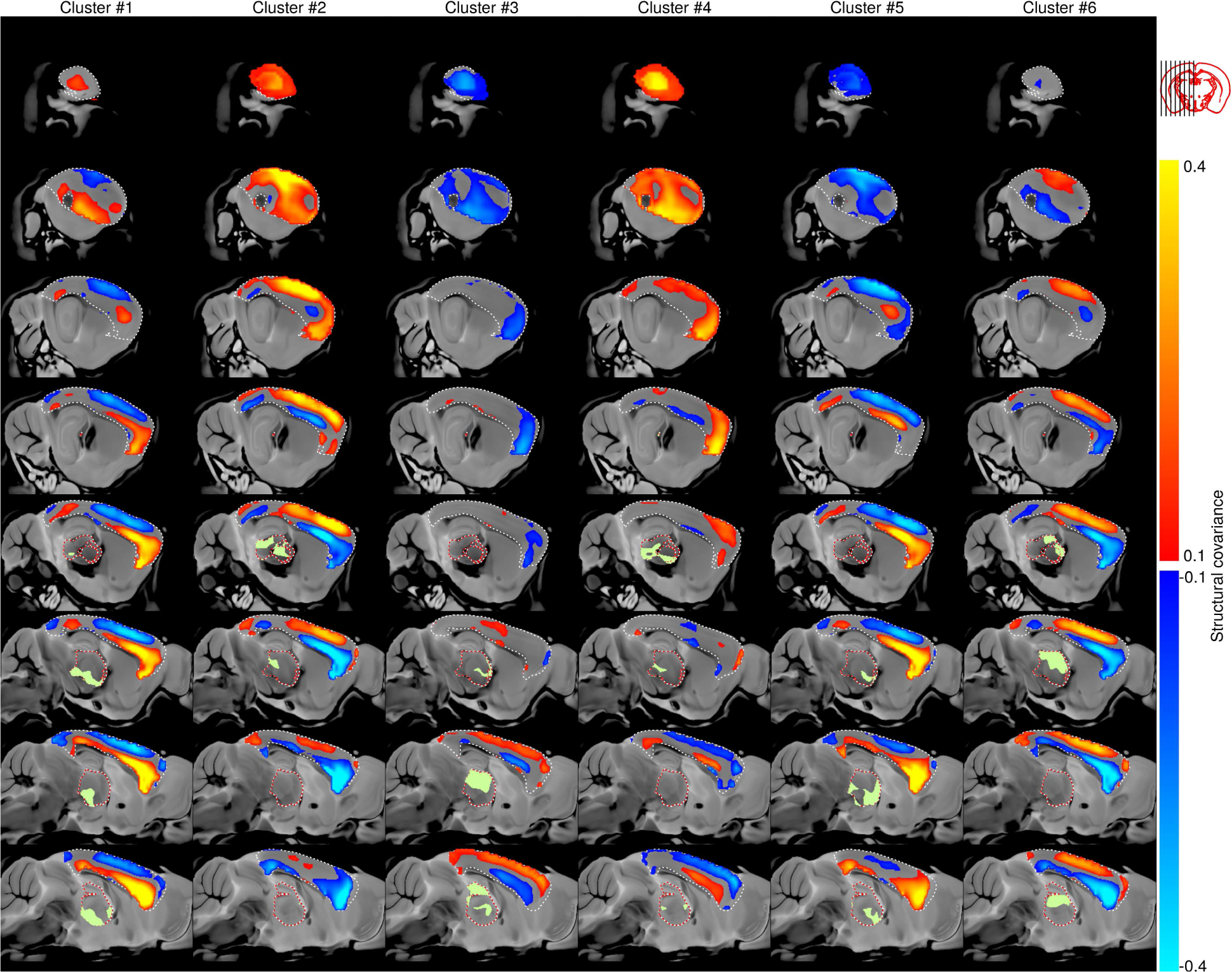
A sagittal view of cortical structural covariance for each of the six clusters. Shown are slices of the average cortical structural covariance map (i.e. *k*-means cluster center) for each of the six clusters of thalamic voxels. Cortical structural covariance is plotted on the study MRI average as the background.

**Figure S8:**
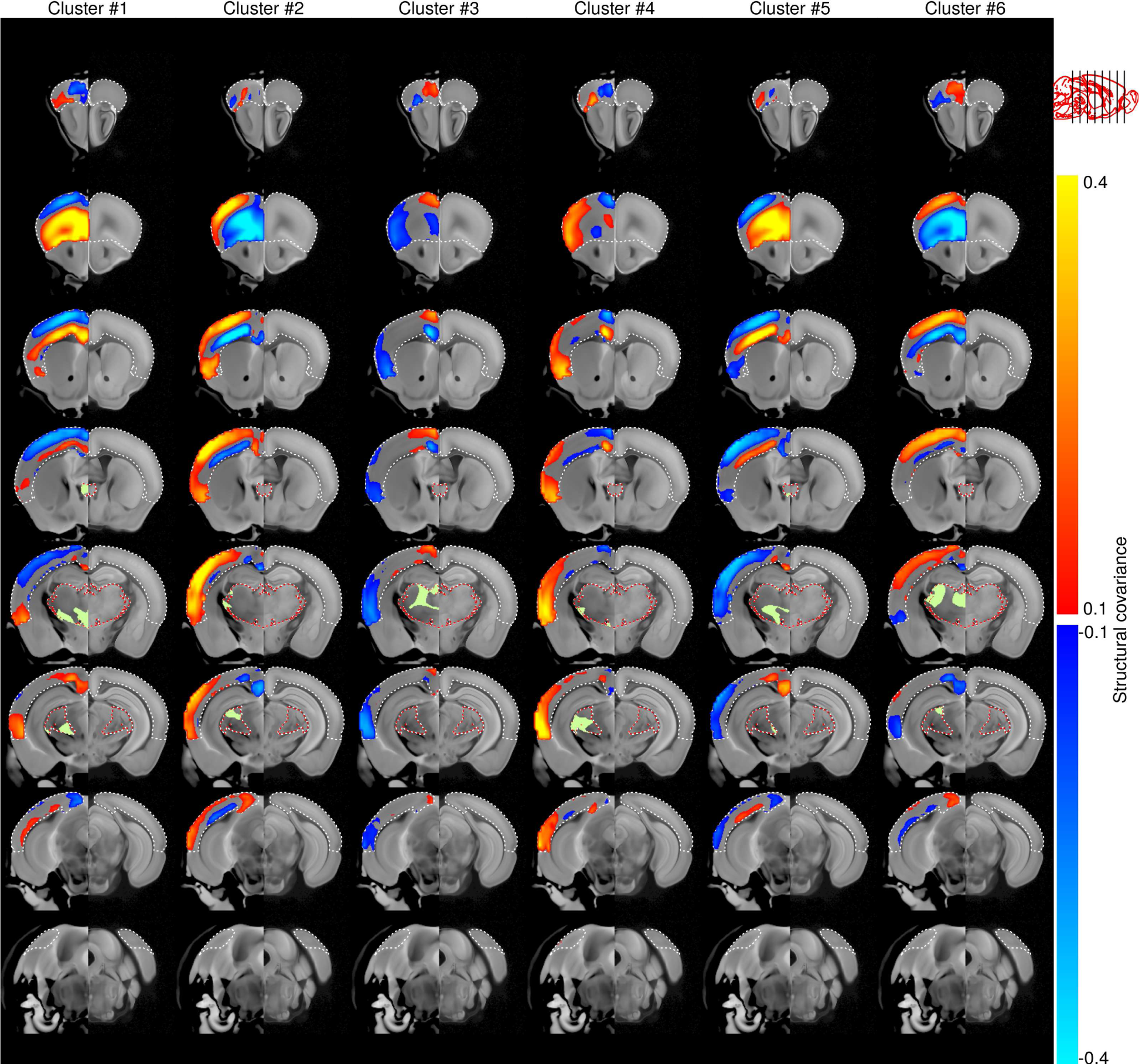
A coronal view of cortical structural covariance for each of the six clusters. Shown are slices of the average cortical structural covariance map (i.e. *k*-means cluster center) for each of the six clusters of thalamic voxels. Cortical structural covariance is plotted on the study MRI average as the background (left hemisphere); the Allen Institute’s CCFv3 template is used as the background in the opposite (right) hemisphere.

**Figure S9:**
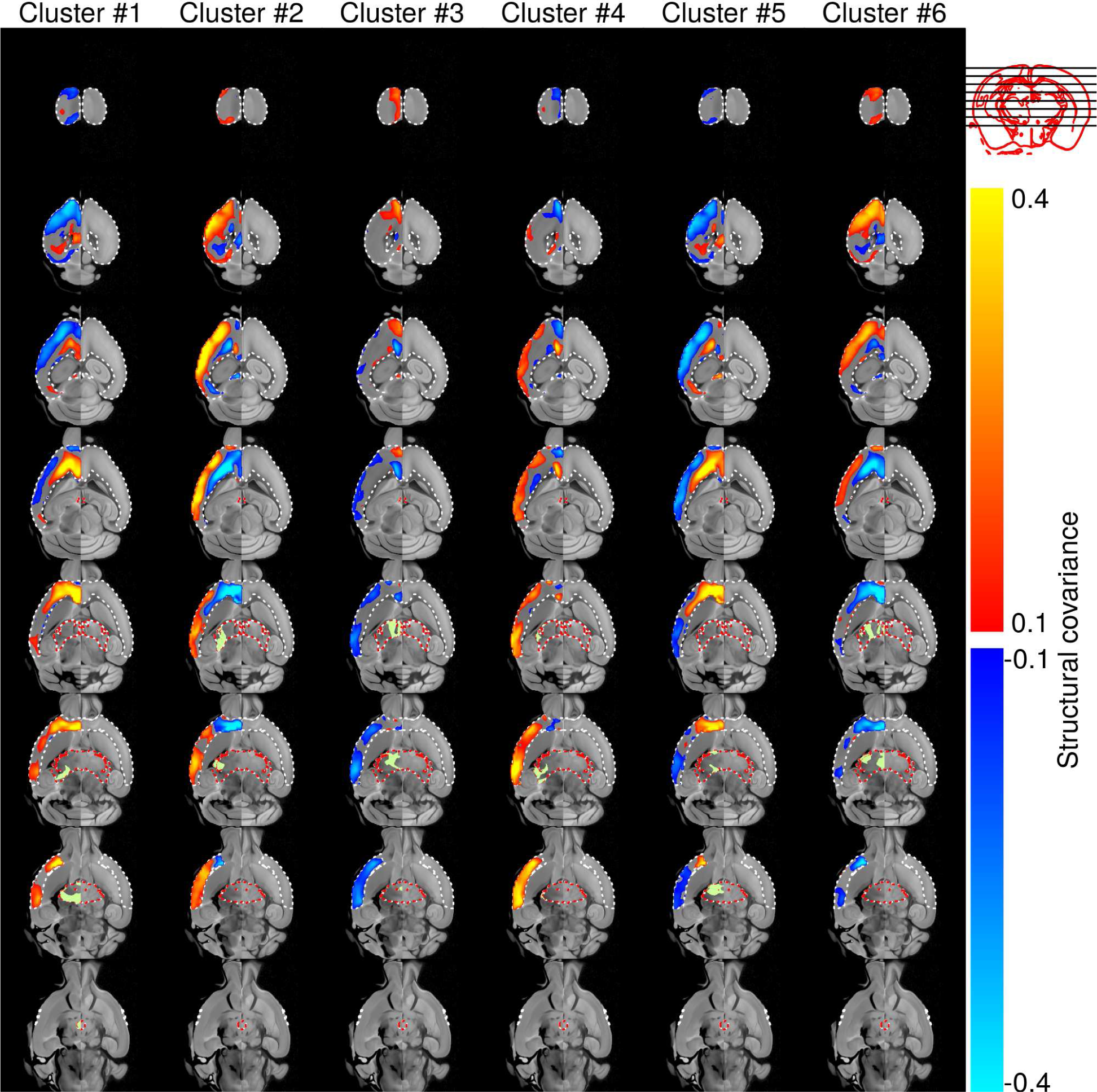
An axial view of cortical structural covariance for each of the six clusters. Shown are slices of the average cortical structural covariance map (i.e. *k*-means cluster center) for each of the six clusters of thalamic voxels. Cortical structural covariance is plotted on the study MRI average as the background (left hemisphere); the Allen Institute’s CCFv3 template is used as the background in the opposite (right) hemisphere.

**Figure S10:**
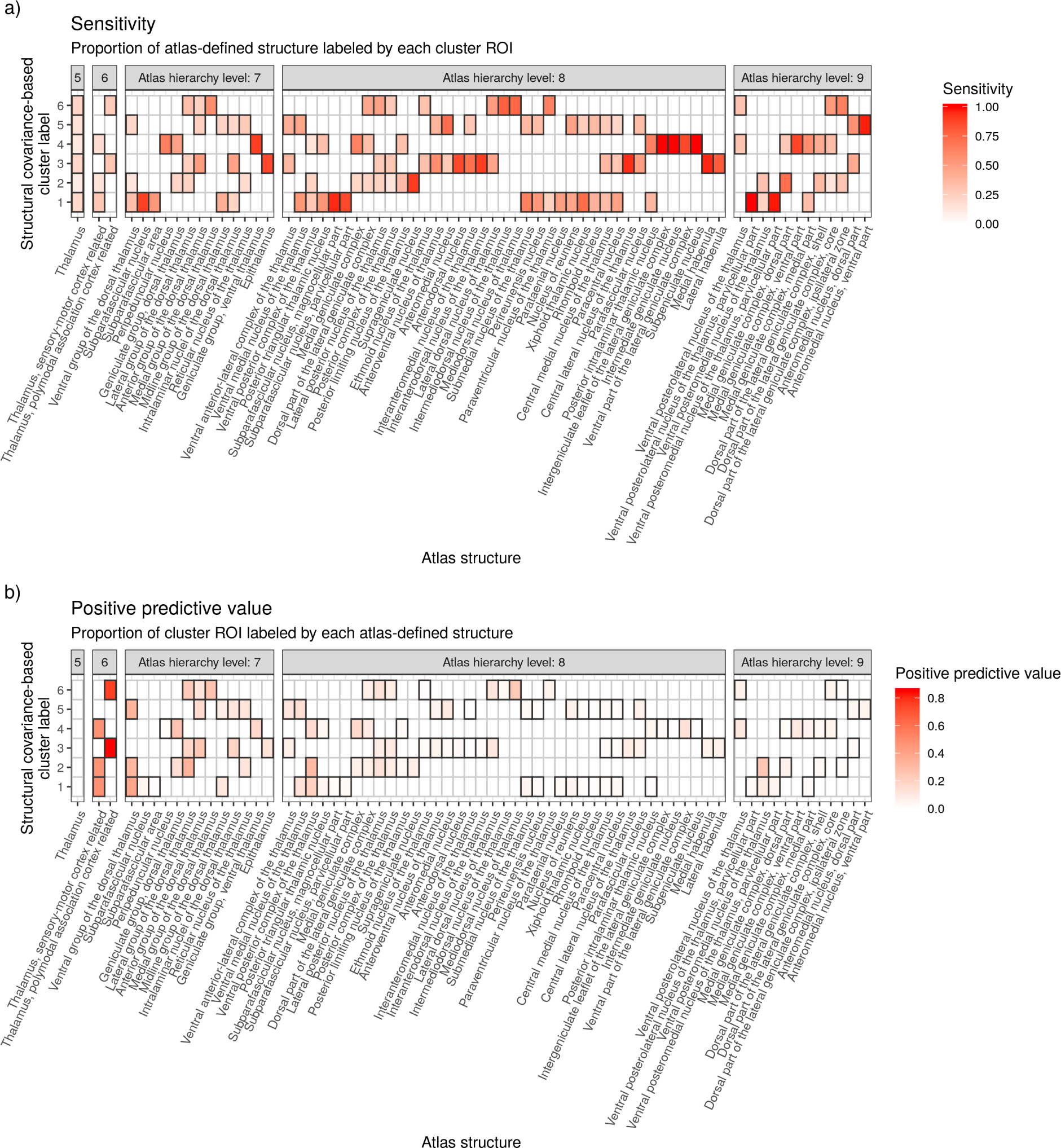
Sensitivity and positive predictive value of structural covariance-based thalamic subdivisions in predicting Allen Mouse Brain Atlas (AMBA) segmentations. For various levels of the AMBA hierarchy, the **a)** sensitivity (proportion that each AMBA segmentation is overlapped by each structural covariance-based cluster) and **b)** positive predictive value (proportion that each covariance-based cluster is overlapped by each AMBA segmentation) is shown. Only statistically significant relationships (assessed via a permutation test and corrected for multiple comparisons) are highlighted.

**Figure S11:**
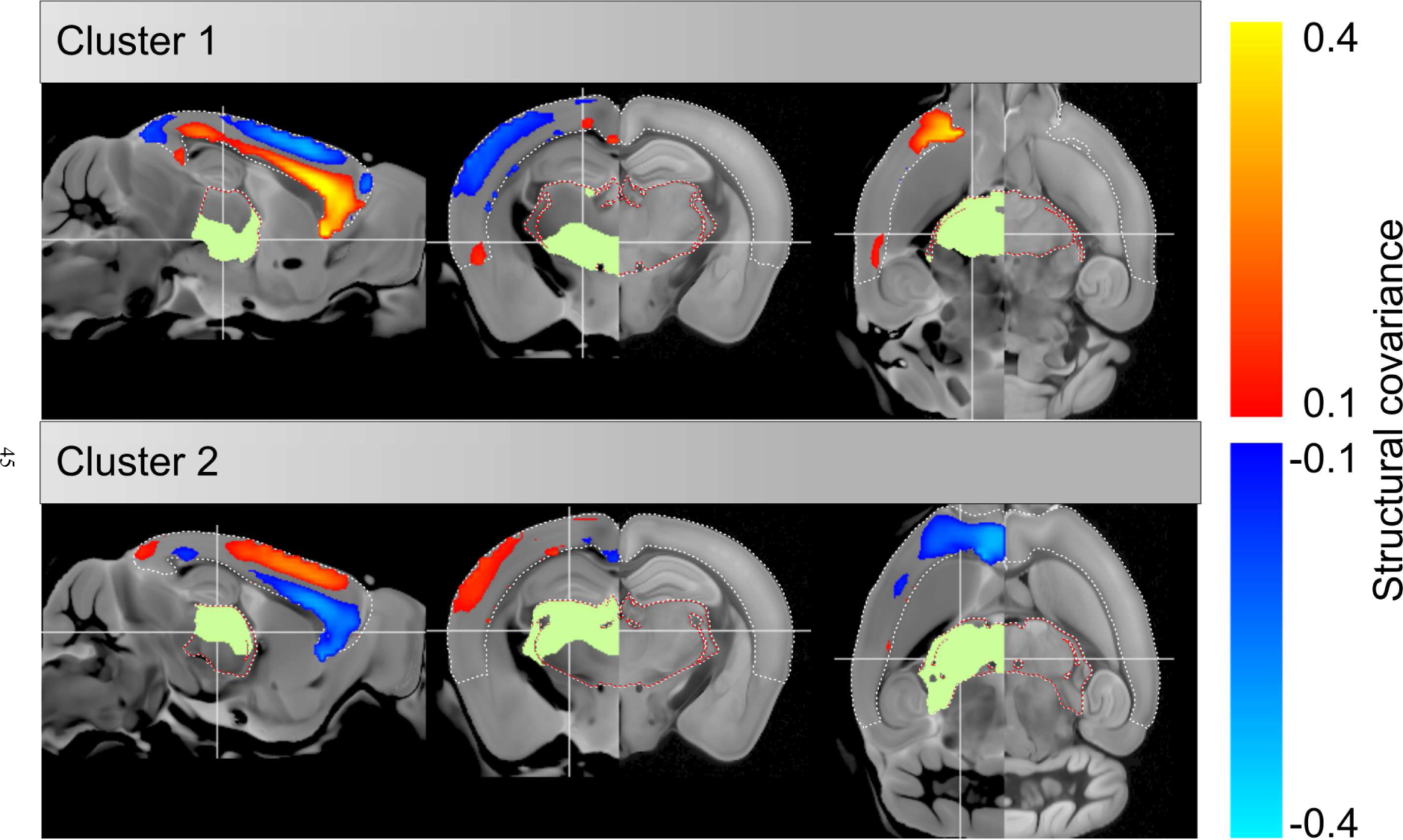
Cortical structural covariance maps associated with structural covariance-based subdivision of the thalamus into two clusters. Two panels that correspond to each cluster show both the thalamic subdivision and associated average cortical structural covariance map, plotted in sagittal, coronal, and axial views.

**Figure S12:**
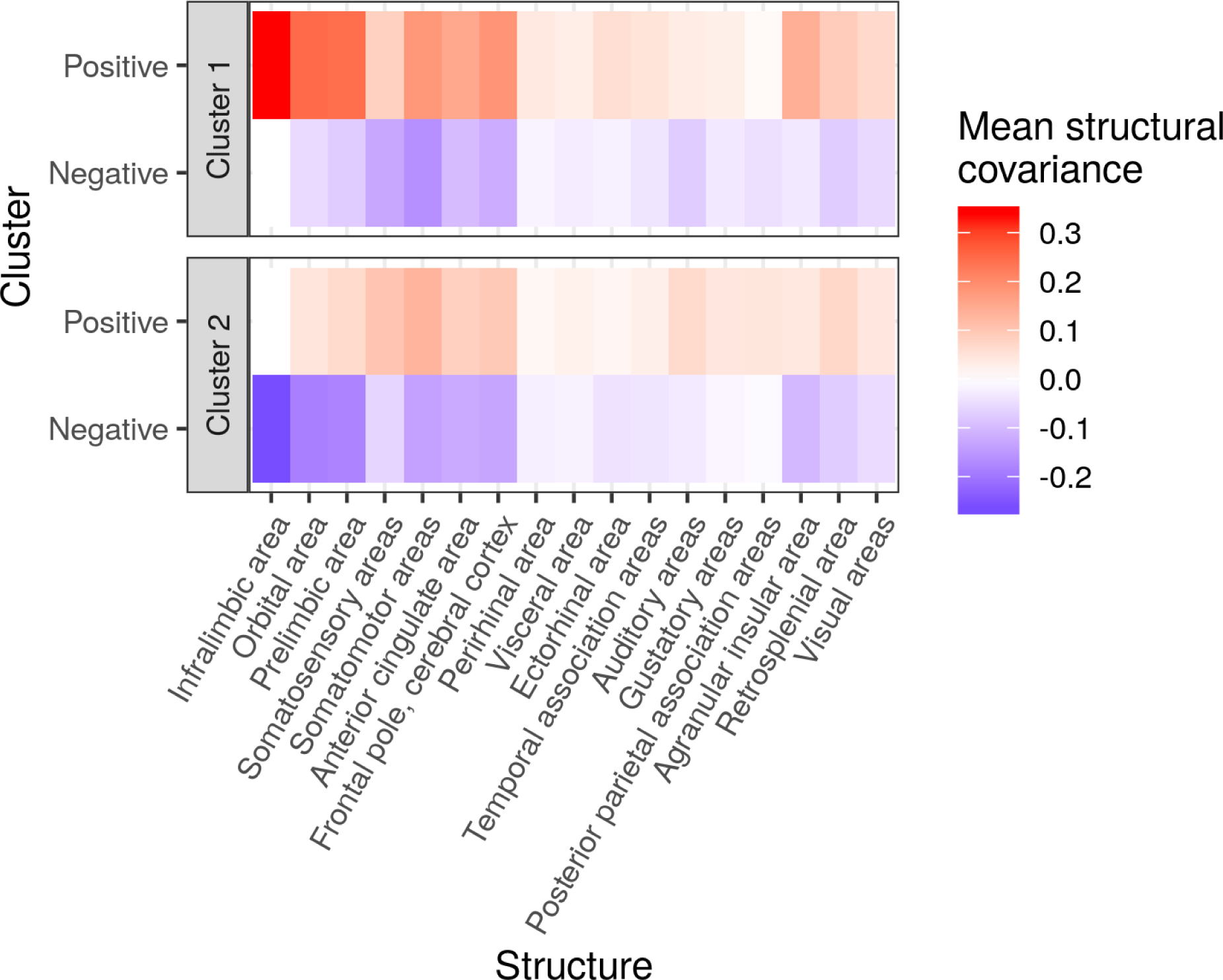
Mean cortical structural covariance for each of the two clusters, summarized under cortical regions. Cortical regions are from the seventh level of the Allen Mouse Brain Atlas hierarchy. For each cluster, mean positive and negative structural covariance under each cortical region are separately calculated by ignoring values of the opposite sign.

**Figure S13:**
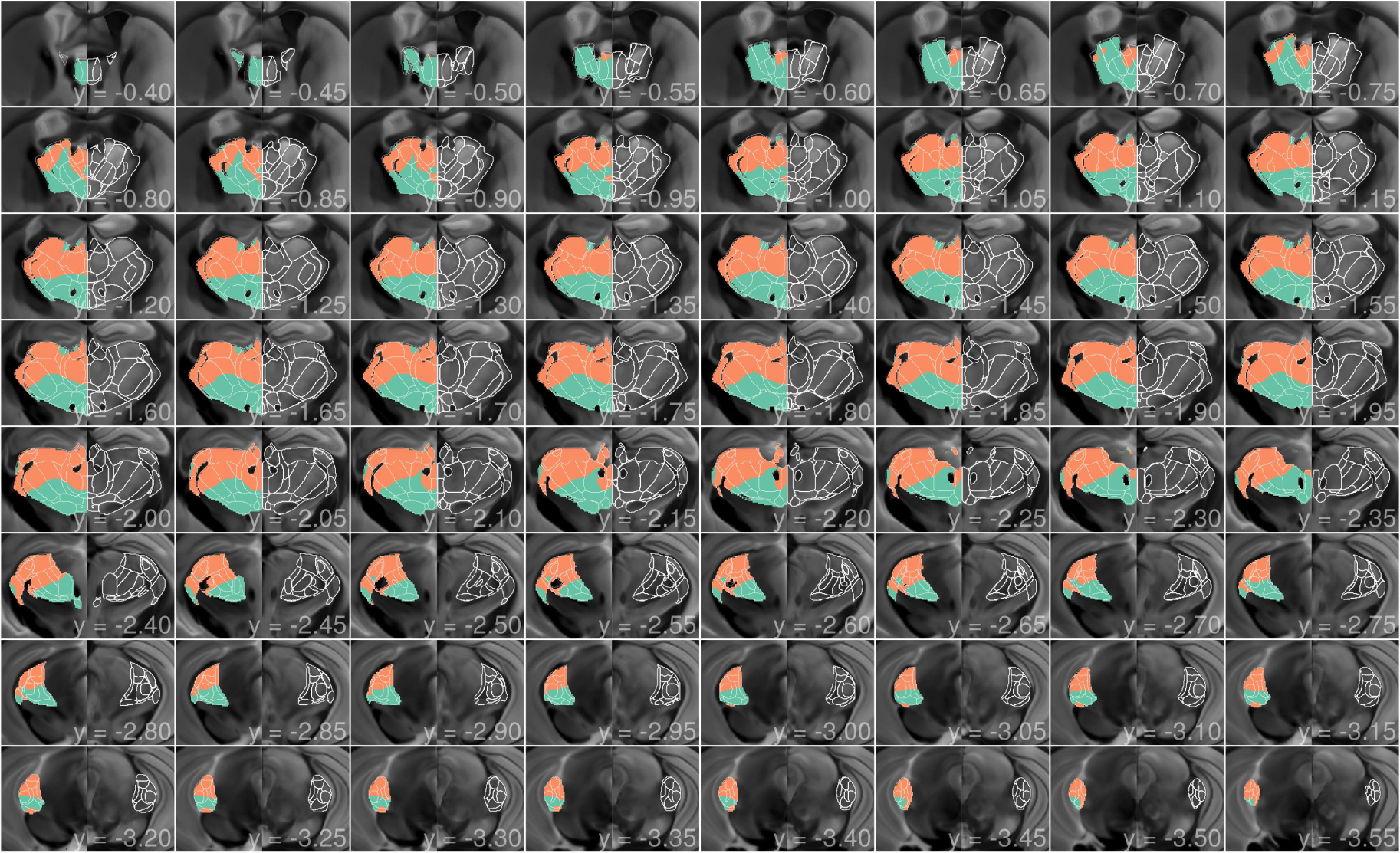
Structural covariance-based subdivision of the thalamus into two clusters. A panel of coronal slices show the thalamus from the anterior to posterior end. Within each slice, covariance-based subdivisions are shown in the left hemisphere on the study MRI average as the background, and contralateral to this, thalamic anatomy is visualized via the CCFv3 template in the right hemisphere. Allen Mouse Brain Atlas segmentations of thalamic nuclei are represented via boundary contours in both hemispheres.

**Figure S14:**
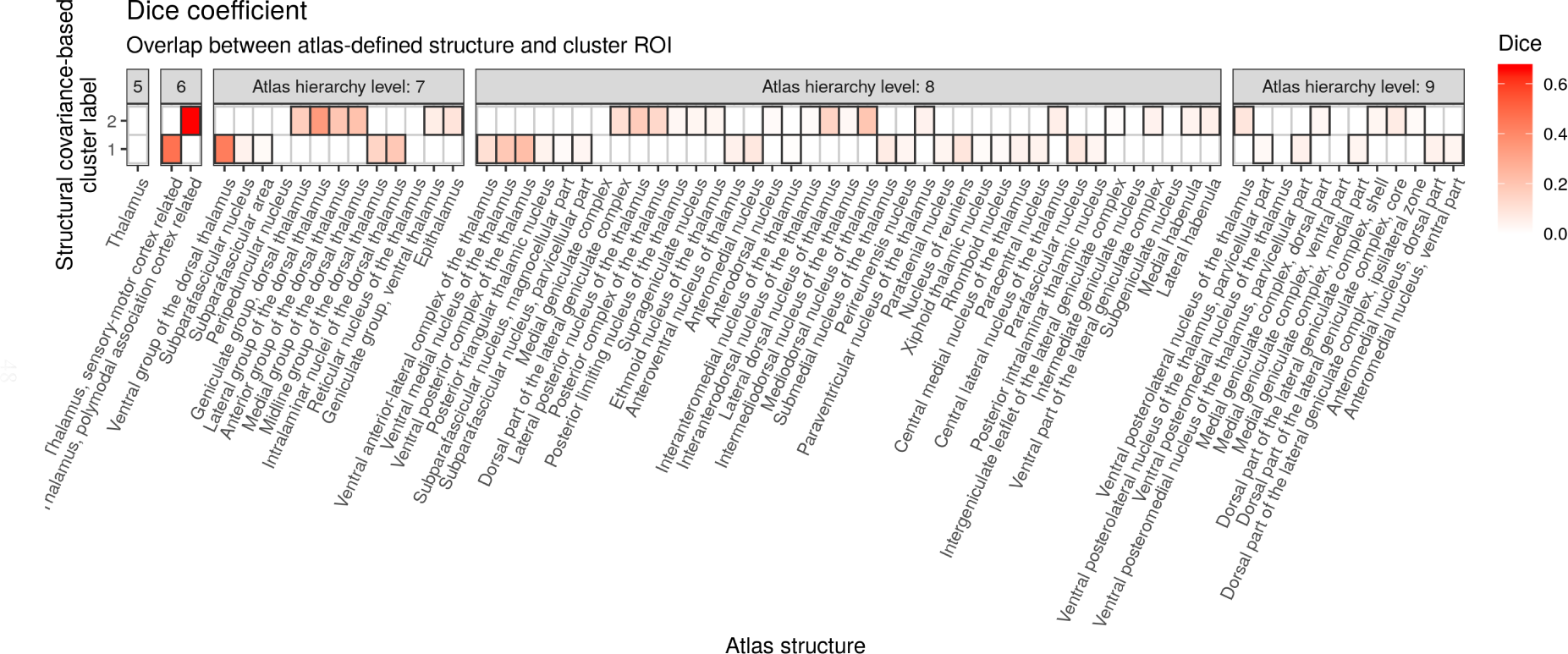
Overlap between two structural covariance-based thalamic subdivisions and the Allen Mouse Brain Atlas (AMBA) segmentations of thalamic nuclei, as quantified by the Dice coefficient. For various levels of the AMBA hierarchy, statistically significant (after controlling for the family-wise error rate) Dice coefficients are plotted.

**Figure S15:**
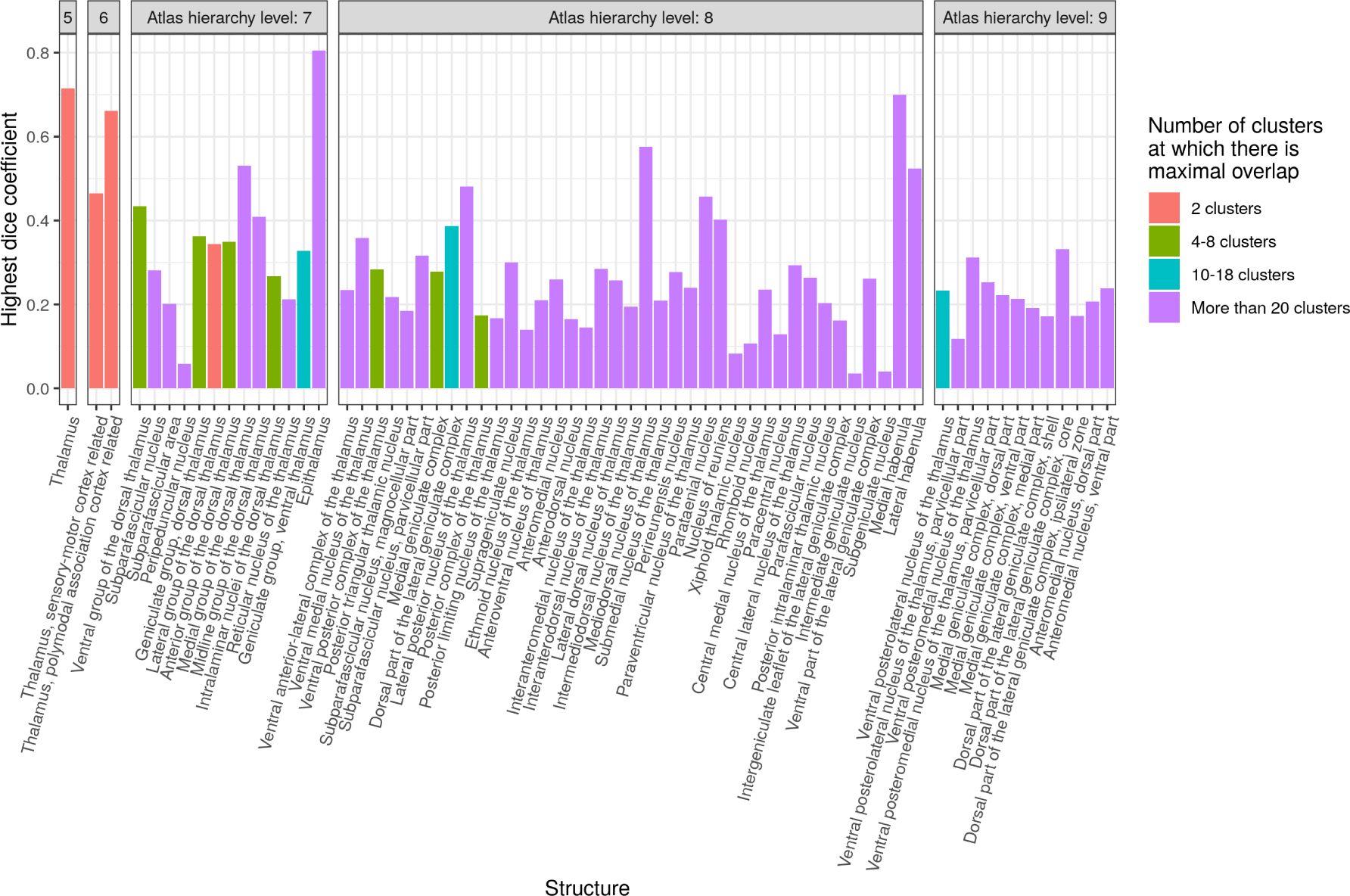
Maximum overlap between Allen Mouse Brain Atlas (AMBA) segmentations of thalamic nuclei and thalamocortical structural covariance-based clusters, for 2 to 42 clusters. For each AMBA segmentation, the maximum overlap (Dice coefficient) with thalamocortical structural covariance-based clusters obtained from a clustering into 2 to 42 clusters is shown. Bar colours indicate how many clusters are required for optimal overlap.

## Footnotes

1 We previously described structural covariance patterns constructed in a group of 153 wildtype mice in Yee et al. (2018). While the sample size reported here is almost identical, this is a different dataset that shares only 10 images with the previously described dataset. The inclusion of the 154 images reported here followed more stringent criteria that minimized age and scan sequence variation; these 154 mice were all young adults at the time of perfusion, and images of their brains were acquired using an identical *T*_2_-weighted fast-spin echo protocol (most recently implemented in-house), unlike the previous dataset which pooled images scanned using slightly different pulse sequences.

## References

Alexander-Bloch, A., Giedd, J. N., & Bullmore, E. (2013a). Imaging structural co-variance between human brain regions. Nature Reviews Neuroscience, 14, 322–336.

Alexander-Bloch, A., Raznahan, A., Bullmore, E., & Giedd, J. (2013b). The convergence of maturational change and structural covariance in human cortical networks. The Journal of Neuroscience, 33, 2889–2899.

Andersen, S. L. (2003). Trajectories of brain development: point of vulnerability or window of opportunity? Neuroscience & Biobehavioral Reviews, 27, 3–18.

Andrews, T. J., Halpern, S. D., & Purves, D. (1997). Correlated size variations in human visual cortex, lateral geniculate nucleus, and optic tract. The Journal of Neuroscience, 17, 2859–2868.

Avants, B. B., Epstein, C. L., Grossman, M., & Gee, J. C. (2008). Symmetric diffeomorphic image registration with cross-correlation: evaluating automated labeling of elderly and neurodegenerative brain. Medical Image Analysis, 12, 26–41.

Avants, B. B., Tustison, N. J., Song, G., Cook, P. A., Klein, A., & Gee, J. C. (2011). A reproducible evaluation of ANTs similarity metric performance in brain image registration. NeuroImage, 54, 2033–2044.

Behrens, T. E., Johansen-Berg, H., Woolrich, M., Smith, S., Wheeler-Kingshott, C., Boulby, P., Barker, G., Sillery, E., Sheehan, K., Ciccarelli, O. et al. (2003). Non-invasive mapping of connections between human thalamus and cortex using diffusion imaging. Nature Neuroscience, 6, 750–757.

Benjamini, Y., & Hochberg, Y. (1995). Controlling the false discovery rate: a practical and powerful approach to multiple testing. Journal of the Royal Statistical Society. Series B (Methodological), (pp. 289–300).

Bernhardt, B. C., Smallwood, J., Keilholz, S., & Margulies, D. S. (2022). Gradients in brain organization. NeuroImage, 251, 118987.

Bogado Lopes, J., Senko, A. N., Bahnsen, K., Geisler, D., Kim, E., Bernanos, M., Cash, D., Ehrlich, S., Vernon, A. C., & Kempermann, G. (2023). Individual behavioral trajectories shape whole-brain connectivity in mice. eLife, 12, e80379.

Bohbot, V. D., Lerch, J., Thorndycraft, B., Iaria, G., & Zijdenbos, A. P. (2007). Gray matter differences correlate with spontaneous strategies in a human virtual navigation task. The Journal of Neuroscience, 27, 10078–10083.

Boretius, S., Kasper, L., Tammer, R., Michaelis, T., & Frahm, J. (2009). MRI of cellular layers in mouse brain in vivo. NeuroImage, 47, 1252–1260.

Chakravarty, M. M., Steadman, P., Van Eede, M. C., Calcott, R. D., Gu, V., Shaw, P., Raznahan, A., Collins, D. L., & Lerch, J. P. (2013). Performing label-fusion-based segmentation using multiple automatically generated templates. Human Brain Mapping, 34, 2635–2654.

Cohen, M. X., Lombardo, M. V., & Blumenfeld, R. S. (2008). Covariance-based subdivision of the human striatum using t1-weighted mri. European Journal of Neuroscience, 27, 1534–1546.

Collins, D. L., Neelin, P., Peters, T. M., & Evans, A. C. (1994). Automatic 3D intersubject registration of MR volumetric data in standardized Talairach space. Journal of Computer Assisted Tomography, 18, 192–205.

Dazai, J., Spring, S., Cahill, L. S., & Henkelman, R. M. (2011). Multiple-mouse neuroanatomical magnetic resonance imaging. Journal of Visualized Experiments, (p. e2497).

Eickhoff, S. B., Yeo, B., & Genon, S. (2018). Imaging-based parcellations of the human brain. Nature Reviews Neuroscience, 19, 672–686.

Elkan, C. (2003). Using the triangle inequality to accelerate k-means. In Proceedings of the 20th international conference on Machine Learning (ICML-03) (pp. 147–153).

Ellegood, J., Anagnostou, E., Babineau, B., Crawley, J., Lin, L., Genestine, M., DiCicco-Bloom, E., Lai, J., Foster, J., Penagarikano, O. et al. (2015). Clustering autism: using neuroanatomical differences in 26 mouse models to gain insight into the heterogeneity. Molecular Psychiatry, 20, 118–125.

Evans, A. C. (2013). Networks of anatomical covariance. NeuroImage, 80, 489–504.

Eyler, L. T., Chen, C.-H., Panizzon, M. S., Fennema-Notestine, C., Neale, M. C., Jak, A., Jernigan, T. L., Fischl, B., Franz, C. E., Lyons, M. J. et al. (2012). A comparison of heritability maps of cortical surface area and thickness and the influence of adjustment for whole brain measures: a magnetic resonance imaging twin study. Twin Research and Human Genetics, 15, 304–314.

Fox, M. D., Zhang, D., Snyder, A. Z., & Raichle, M. E. (2009). The global signal and observed anticorrelated resting state brain networks. Journal of Neurophysiology, 101, 3270–3283.

Fracasso, A., van Veluw, S. J., Visser, F., Luijten, P. R., Spliet, W., Zwanenburg, J. J., Dumoulin, S. O., & Petridou, N. (2016). Lines of Baillarger in vivo and ex vivo: myelin contrast across lamina at 7 T MRI and histology. NeuroImage, 133, 163–175.

French, L., & Pavlidis, P. (2011). Relationships between gene expression and brain wiring in the adult rodent brain. PLOS Computational Biology, 7, e1001049.

Friedel, M., van Eede, M. C., Pipitone, J., Chakravarty, M. M., & Lerch, J. P. (2014). Pydpiper: a flexible toolkit for constructing novel registration pipelines. Frontiers in Neuroinformatics, 8, 67.

Fukunaga, M., Li, T.-Q., van Gelderen, P., de Zwart, J. A., Shmueli, K., Yao, B., Lee, J., Maric, D., Aronova, M. A., Zhang, G. et al. (2010). Layer-specific variation of iron content in cerebral cortex as a source of MRI contrast. Proceedings of the National Academy of Sciences, 107, 3834–3839.

Gong, G., He, Y., Chen, Z. J., & Evans, A. C. (2012). Convergence and divergence of thickness correlations with diffusion connections across the human cerebral cortex. NeuroImage, 59, 1239–1248.

Guillery, R., & Sherman, S. M. (2002). Thalamic relay functions and their role in corticocortical communication: generalizations from the visual system. Neuron, 33, 163–175.

Harris, J. A., Mihalas, S., Hirokawa, K. E., Whitesell, J. D., Choi, H., Bernard, A., Bohn, P., Caldejon, S., Casal, L., Cho, A. et al. (2019). Hierarchical organization of cortical and thalamic connectivity. Nature, 575, 195–202.

Hashikawa, T., Molinari, M., Rausell, E., & Jones, E. (1995). Patchy and laminar terminations of medial geniculate axons in monkey auditory cortex. Journal of Comparative Neurology, 362, 195–208.

Hendrickson, A., Wilson, J., & Ogren, M. (1978). The neuroanatomical organization of pathways between the dorsal lateral geniculate nucleus and visual cortex in Old World and New World primates. Journal of Comparative Neurology, 182, 123–136.

Hollander, H. (1972). Autoradiographic evidence for a projection from the striate cortex to the dorsal part of the lateral geniculate nucleus in the cat. Brain Research, 41, 464.

Jha, S. C., Xia, K., Schmitt, J. E., Ahn, M., Girault, J. B., Murphy, V. A., Li, G., Wang, L., Shen, D., Zou, F. et al. (2018). Genetic influences on neonatal cortical thickness and surface area. Human Brain Mapping, 39, 4998–5013.

Ji, B., Li, Z., Li, K., Li, L., Langley, J., Shen, H., Nie, S., Zhang, R., & Hu, X. (2016). Dynamic thalamus parcellation from resting-state fMRI data. Human Brain Mapping, 37, 954–967.

Johansen-Berg, H., Behrens, T. E., Sillery, E., Ciccarelli, O., Thompson, A. J., Smith, S. M., & Matthews, P. M. (2005). Functional–anatomical validation and individual variation of diffusion tractography-based segmentation of the human thalamus. Cerebral Cortex, 15, 31–39.

Jones, E. (1998). Viewpoint: the core and matrix of thalamic organization. Neuroscience, 85, 331–345.

Jones, E., & Powell, T. (1970). Connexions of the somatic sensory cortex of the rhesus monkey: Iii.—thalamic connexions. Brain, 93, 37–56.

Jones, E., Wise, S., & Coulter, J. (1979). Differential thalamic relationships of sensory-motor and parietal cortical fields in monkeys. Journal of Comparative Neurology, 183, 833–881.

Jones, E. G. (1985). The thalamus. Springer Science & Business Media.

Jones, S. E., Buchbinder, B. R., & Aharon, I. (2000). Three-dimensional mapping of cortical thickness using Laplace’s equation. Human Brain Mapping, 11, 12–32.

Joshi, A. A., Lepore, N., Joshi, S. H., Lee, A. D., Barysheva, M., Stein, J. L., McMahon, K. L., Johnson, K., de Zubicaray, G. I., Martin, N. G. et al. (2011). The contribution of genes to cortical thickness and volume. NeuroReport, 22, 101.

Kelly, C., Toro, R., Di Martino, A., Cox, C. L., Bellec, P., Castellanos, F. X., & Milham, M. P. (2012). A convergent functional architecture of the insula emerges across imaging modalities. NeuroImage, 61, 1129–1142.

Krubitzer, L., & Huffman, K. J. (2000). Arealization of the neocortex in mammals: genetic and epigenetic contributions to the phenotype. Brain, Behavior and Evolution, 55, 322–335.

Lambert, C., Simon, H., Colman, J., & Barrick, T. R. (2017). Defining thalamic nuclei and topographic connectivity gradients in vivo. NeuroImage, 158, 466–479.

Lee, J., Ehlers, C., Crews, F., Niethammer, M., Budin, F., Paniagua, B., Sulik, K., Johns, J., Styner, M., & Oguz, I. (2011). Automatic cortical thickness analysis on rodent brain. In Medical Imaging 2011: Image Processing (p. 796248). International Society for Optics and Photonics volume 7962.

Lerch, J., Hammill, C., van Eede, M., & Cassel, D. (2017). RMINC: Statistical tools for Medical Imaging NetCDF (MINC) files. URL: http://mouse-imaging-centre.github.io/{RMINC} R package version 1.5.2.1.

Lerch, J. P., Carroll, J. B., Dorr, A., Spring, S., Evans, A. C., Hayden, M. R., Sled, J. G., & Henkelman, R. M. (2008). Cortical thickness measured from MRI in the YAC128 mouse model of Huntington’s disease. NeuroImage, 41, 243–251.

Lerch, J. P., Sled, J. G., & Henkelman, R. M. (2011). MRI phenotyping of genetically altered mice. Methods in Molecular Biology, 711, 349–361.

Lerch, J. P., Worsley, K., Shaw, W. P., Greenstein, D. K., Lenroot, R. K., Giedd, J., & Evans, A. C. (2006). Mapping anatomical correlations across cerebral cortex (MACACC) using cortical thickness from MRI. NeuroImage, 31, 993–1003.

Liang, X., Wang, J., Yan, C., Shu, N., Xu, K., Gong, G., & He, Y. (2012). Effects of different correlation metrics and preprocessing factors on small-world brain functional networks: a resting-state functional MRI study. PLoS One, 7, e32766.

Long, J., Lu, F., Guo, X., Pang, Y., Yang, S., Chen, H., & He, B. (2021). Parcellation of the thalamus by using a dual-segment method based on resting-state functional connectivity: An application on autism spectrum disorder. Neuroscience Letters, 742, 135518.

López-Bendito, G., & Molnár, Z. (2003). Thalamocortical development: how are we going to get there? Nature Reviews Neuroscience, 4, 276–289.

Magnotta, V. A., Gold, S., Andreasen, N. C., Ehrhardt, J. C., & Yuh, W. T. (2000). Visualization of subthalamic nuclei with cortex attenuated inversion recovery MR imaging. NeuroImage, 11, 341–346.

Masouleh, S. K., Plachti, A., Hoffstaedter, F., Eickhoff, S., & Genon, S. (2020). Characterizing the gradients of structural covariance in the human hippocampus. NeuroImage, 218, 116972.

Mechelli, A., Friston, K. J., Frackowiak, R. S., & Price, C. J. (2005). Structural covariance in the human cortex. The Journal of Neuroscience, 25, 8303–8310.

Middlebrooks, E. H., Tuna, I. S., Almeida, L., Grewal, S. S., Wong, J., Heckman, M. G., Lesser, E. R., Bredel, M., Foote, K. D., Okun, M. S. et al. (2018). Structural connectivity-based segmentation of the thalamus and prediction of tremor improvement following thalamic deep brain stimulation of the ventral intermediate nucleus. NeuroImage: Clinical, 20, 1266–1273.

Molinari, M., Dell’Anna, M., Rausell, E., Leggio, M., Hashikawa, T., & Jones, E. (1995). Auditory thalamocortical pathways defined in monkeys by calcium-binding protein immunoreactivity. Journal of Comparative Neurology, 362, 171–194.

Müller, E. J., Munn, B., Hearne, L. J., Smith, J. B., Fulcher, B., Arnatkevičiūtė, A., Lurie, D. J., Cocchi, L., & Shine, J. M. (2020). Core and matrix thalamic sub-populations relate to spatio-temporal cortical connectivity gradients. NeuroImage, 222, 117224.

Nieman, B. J., van Eede, M. C., Spring, S., Dazai, J., Henkelman, R. M., & Lerch, J. P. (2018). MRI to assess neurological function. Current Protocols in Mouse Biology, 8, e44.

Oh, S. W., Harris, J. A., Ng, L., Winslow, B., Cain, N., Mihalas, S., Wang, Q., Lau, C., Kuan, L., Henry, A. M. et al. (2014). A mesoscale connectome of the mouse brain. Nature, 508, 207–214.

O’Muircheartaigh, J., Vollmar, C., Traynor, C., Barker, G. J., Kumari, V., Symms, M. R., Thompson, P., Duncan, J. S., Koepp, M. J., & Richardson, M. P. (2011). Clustering probabilistic tractograms using independent component analysis applied to the thalamus. NeuroImage, 54, 2020–2032.

Pagani, M., Bifone, A., & Gozzi, A. (2016). Structural covariance networks in the mouse brain. NeuroImage, 129, 55–63.

Passingham, R. E., Stephan, K. E., & Kötter, R. (2002). The anatomical basis of functional localization in the cortex. Nature Reviews Neuroscience, 3, 606–616.

Pedregosa, F., Varoquaux, G., Gramfort, A., Michel, V., Thirion, B., Grisel, O., Blondel, M., Prettenhofer, P., Weiss, R., Dubourg, V. et al. (2011). Scikit-learn: Machine learning in Python. The Journal of Machine Learning Research, 12, 2825–2830.

Persson, J., Spreng, R. N., Turner, G., Herlitz, A., Morell, A., Stening, E., Wahlund, L.-O., Wikström, J., & Söderlund, H. (2014). Sex differences in volume and structural covariance of the anterior and posterior hippocampus. Neuroimage, 99, 215–225.

Phillips, J. W., Schulmann, A., Hara, E., Winnubst, J., Liu, C., Valakh, V., Wang, L., Shields, B. C., Korff, W., Chandrashekar, J. et al. (2019). A repeated molecular architecture across thalamic pathways. Nature Neuroscience, 22, 1925–1935.

Plachti, A., Eickhoff, S. B., Hoffstaedter, F., Patil, K. R., Laird, A. R., Fox, P. T., Amunts, K., & Genon, S. (2019). Multimodal parcellations and extensive behavioral profiling tackling the hippocampus gradient. Cerebral Cortex, 29, 4595–4612.

R Core Team (2020). R: A language and environment for statistical computing. R Foundation for Statistical Computing Vienna, Austria. URL: https://www.R-project.org/.

Rakic, P. (1988). Specification of cerebral cortical areas. Science, 241, 170–176.

Raznahan, A., Lerch, J. P., Lee, N., Greenstein, D., Wallace, G. L., Stockman, M., Clasen, L., Shaw, P. W., & Giedd, J. N. (2011). Patterns of coordinated anatomical change in human cortical development: a longitudinal neuroimaging study of maturational coupling. Neuron, 72, 873–884.

Richiardi, J., Altmann, A., Milazzo, A.-C., Chang, C., Chakravarty, M. M., Banaschewski, T., Barker, G. J., Bokde, A. L., Bromberg, U., Büchel, C. et al. (2015). Correlated gene expression supports synchronous activity in brain networks. Science, 348, 1241–1244.

Romero-Garcia, R., Whitaker, K. J., Váša, F., Seidlitz, J., Shinn, M., Fonagy, P., Dolan, R. J., Jones, P. B., Goodyer, I. M., Bullmore, E. T. et al. (2018). Structural covariance networks are coupled to expression of genes enriched in supragranular layers of the human cortex. NeuroImage, 171, 256–267.

Sanabria-Diaz, G., Melie-García, L., Iturria-Medina, Y., Alemán-Gómez, Y., Hernández-González, G., Valdés-Urrutia, L., Galán, L., & Valdés-Sosa, P. (2010). Surface area and cortical thickness descriptors reveal different attributes of the structural human brain networks. NeuroImage, 50, 1497–1510.

Scrucca, L., Fop, M., Murphy, T. B., & Raftery, A. E. (2016). mclust 5: clustering, classification and density estimation using Gaussian finite mixture models. The R Journal, 8, 289.

Segall, J. M., Allen, E. A., Jung, R. E., Erhardt, E. B., Arja, S. K., Kiehl, K. A., & Calhoun, V. D. (2012). Correspondence between structure and function in the human brain at rest. Frontiers in Neuroinformatics, 6, 10.

Sherman, S. M. (2007). The thalamus is more than just a relay. Current Opinion in Neurobiology, 17, 417–422.

Sherman, S. M. (2016). Thalamus plays a central role in ongoing cortical functioning. Nature Neuroscience, 19, 533–541.

Sherman, S. M., & Guillery, R. (2002). The role of the thalamus in the flow of information to the cortex. Philosophical Transactions of the Royal Society of London. Series B: Biological Sciences, 357, 1695–1708.

Sled, J. G., Zijdenbos, A. P., & Evans, A. C. (1998). A nonparametric method for automatic correction of intensity nonuniformity in MRI data. IEEE Transactions on Medical Imaging, 17, 87–97.

Spencer Noakes, T. L., Henkelman, R. M., & Nieman, B.J. (2017). Partitioning k-space for cylindrical three-dimensional rapid acquisition with relaxation enhancement imaging in the mouse brain. NMR in Biomedicine, 30, e3802.

Strike, L. T., Hansell, N. K., Couvy-Duchesne, B., Thompson, P. M., de Zubicaray, G. I., McMahon, K. L., & Wright, M. J. (2019). Genetic complexity of cortical structure: differences in genetic and environmental factors influencing cortical surface area and thickness. Cerebral Cortex, 29, 952–962.

Stüber, C., Morawski, M., Schäfer, A., Labadie, C., Wähnert, M., Leuze, C., Streicher, M., Barapatre, N., Reimann, K., Geyer, S. et al. (2014). Myelin and iron concentration in the human brain: a quantitative study of MRI contrast. NeuroImage, 93, 95–106.

Tange, O. (2011). Gnu parallel: The command-line power tool.; login: The USENIX Magazine, 36, 42–47.

Thompson, C. L., Ng, L., Menon, V., Martinez, S., Lee, C.-K., Glattfelder, K., Sunkin, S. M., Henry, A., Lau, C., Dang, C. et al. (2014). A high-resolution spatiotemporal atlas of gene expression of the developing mouse brain. Neuron, 83, 309–323.

Toulmin, H., Beckmann, C. F., O’Muircheartaigh, J., Ball, G., Nongena, P., Makropoulos, A., Ederies, A., Counsell, S. J., Kennea, N., Arichi, T. et al. (2015). Specialization and integration of functional thalamocortical connectivity in the human infant. Proceedings of the National Academy of Sciences, 112, 6485–6490.

Traynor, C., Heckemann, R. A., Hammers, A., O’Muircheartaigh, J., Crum, W. R., Barker, G. J., & Richardson, M. P. (2010). Reproducibility of thalamic segmentation based on probabilistic tractography. NeuroImage, 52, 69–85.

Vincent, R. D., Neelin, P., Khalili-Mahani, N., Janke, A. L., Fonov, V. S., Robbins, S. M., Baghdadi, L., Lerch, J., Sled, J. G., Adalat, R. et al. (2016). Minc 2.0: a flexible format for multi-modal images. Frontiers in Neuroinformatics, 10, 35.

Wang, Q., Ding, S.-L., Li, Y., Royall, J., Feng, D., Lesnar, P., Graddis, N., Naeemi, M., Facer, B., Ho, A. et al. (2020). The Allen Mouse Brain Common Coordinate Framework: a 3D reference atlas. Cell,.

Weissenbacher, A., Kasess, C., Gerstl, F., Lanzenberger, R., Moser, E., & Windischberger, C. (2009). Correlations and anticorrelations in resting-state functional connectivity MRI: a quantitative comparison of preprocessing strategies. NeuroImage, 47, 1408–1416.

Winkler, A. M., Kochunov, P., Blangero, J., Almasy, L., Zilles, K., Fox, P. T., Duggirala, R., & Glahn, D. C. (2010). Cortical thickness or grey matter volume? the importance of selecting the phenotype for imaging genetics studies. NeuroImage, 53, 1135–1146.

Yang, J.-J., Kwon, H., & Lee, J.-M. (2016). Complementary characteristics of correlation patterns in morphometric correlation networks of cortical thickness, surface area, and gray matter volume. Scientific Reports, 6, 26682.

Yee, Y., Fernandes, D. J., French, L., Ellegood, J., Cahill, L. S., Vousden, D. A., Noakes, L. S., Scholz, J., van Eede, M. C., Nieman, B. J. et al. (2018). Structural covariance of brain region volumes is associated with both structural connectivity and transcriptomic similarity. NeuroImage, 179, 357–372.

Yuan, R., Di, X., Taylor, P. A., Gohel, S., Tsai, Y.-H., & Biswal, B. B. (2016). Functional topography of the thalamocortical system in human. Brain Structure and Function, 221, 1971–1984.

Zalesky, A., Fornito, A., & Bullmore, E. (2012). On the use of correlation as a measure of network connectivity. NeuroImage, 60, 2096–2106.

Zhang, D., Snyder, A. Z., Fox, M. D., Sansbury, M. W., Shimony, J. S., & Raichle, M. E. (2008). Intrinsic functional relations between human cerebral cortex and thalamus. Journal of Neurophysiology, 100, 1740–1748.

